# Rescue of transcriptional pausing in metabolic genes jumpstarts *Salmonella* antioxidant defenses

**DOI:** 10.1101/2022.06.09.495436

**Authors:** Sashi Kant, James Karl A Till, Lin Liu, Ju-Sim Kim, Andres Vazquez-Torres

## Abstract

Detoxification, scavenging and repair systems embody the archetypical antioxidant defenses of prokaryotic and eukaryotic cells^1–3^. Metabolic rewiring is an emergent aspect in the adaptation of bacteria to oxidative stress^4, 5^. Evolutionarily diverse bacteria combat the toxicity of reactive oxygen species by actively engaging the stringent response^6–8^, a metabolic program activated at the level of transcription initiation via guanosine tetraphosphate and the α-helical DksA protein^9^. Studies herein with *Salmonella* demonstrate that the interactions of structurally related, but functionally unique, α-helical Gre factors with the secondary channel of RNA polymerase elicit the expression of metabolic signatures that are associated with resistance to oxidative killing. Gre proteins resolve pauses in ternary elongation complexes of Embden-Meyerhof-Parnas (EMP) glycolysis and aerobic respiration genes. The Gre-directed utilization of glucose in overflow and aerobic metabolism satisfies the energetic and redox demands of *Salmonella*, while preventing the occurrence of amino acid bradytrophies. Moreover, the simultaneous utilization of lower glycolysis, the methylglyoxal pathway and the electron transport chain curtails the noxious coexistence of reductive and electrophilic stress. The resolution of transcriptional pauses in EMP glycolysis and aerobic respiration genes by Gre factors safeguards *Salmonella* from the cytotoxicity of phagocyte NADPH oxidase in the innate host response. Control of transcriptional elongation represents a pivotal breakpoint in the regulation of metabolic programs underlying bacterial pathogenesis.

## INTRODUCTION

Oxidative stress is one of the most potent effectors of the innate immune system^10,11,12^. Reactive oxygen species (ROS) generated by NADPH oxidase (NOX2) in the respiratory burst of phagocytic cells exert potent anti-*Salmonella* activity^13^, damaging nucleic acids, amino acid residues and metal prosthetic groups^14^. Considerable attention has been paid to the antioxidant defenses that counteract the tremendous selective pressures of respiratory burst products. Effectors of the *Salmonella* pathogenicity island-2 (SPI-2) type III secretion system divert NOX2-containing vesicles away from phagosomes^15^. Iron-sequestering proteins prevent formation of genotoxic ferryl and hydroxyl radicals in the Fe^2+^-mediated reduction of hydrogen peroxide (H_2_O_2_)^16^. ROS that reach intracellular *Salmonella* are detoxified by periplasmic superoxide dismutases^17^, or are scavenged by the low-molecular weight thiol glutathione (GSH)^18^. Despite all of these protective mechanisms, ROS are still able to damage the genome and proteome, necessitating restoration by DNA repair systems and thioredoxin^19, 20^. In addition to these classical antioxidant defenses, recent investigations have revealed that metabolic reprogramming is a potent but little understood mechanism in the adaptation of bacteria to the vigorous antimicrobial activity of ROS engendered in the innate host response. *Salmonella* sustaining oxidative stress favor glycolysis, fermentation and the reductive tricarboxylic acid cycle, while slowing down aerobic respiration and oxidative phosphorylation^5^. By doing so, this facultative intracellular pathogen not only diminishes the adventitious generation of superoxide anion in the electron transport chain (ETC), but also allows for the vectorial delivery of electrons by the Dsb thiol-disulfide exchange system into oxidized quinones^5^. Shifting metabolism from oxidative to substrate-level phosphorylation facilitates repair of oxidized periplasmic proteins, while continuing to cover energetic and redox balancing needs^5^.

By recruiting RNA polymerase to the promoters of target genes, transcription factors such as Fnr, PhoP or SsrB as well as alternative σ^S^ and σ^E^ sigma factors activate expression of classical antioxidant defenses^21–25^. *Salmonella* have also harnessed the regulatory activity that the nucleotide alarmone guanosine tetraphosphate (ppGpp) and the small protein DksA exert on RNA polymerase to adapt against NOX2-dependent cytotoxicity^6, 9^. DksA accesses the catalytic site of RNA polymerase via its secondary channel. Two aspartic acid residues at the tip of the coiled-coil domain of DksA clash with the bridge helix of RNA polymerase, collapsing open complexes in the transcription bubble^26^. Direct interactions of DksA with RNA polymerase boost *Salmonella’*s antioxidant defenses by both controlling redox balance and stimulating SPI-2 gene transcription^6, 27^. The transcriptional control of antioxidant defenses exercised by the stringent response illustrates the crosstalk between metabolism and the adaptation to oxidative stress.

The secondary channel of RNA polymerase also accommodates Gre factors, which share similarities with DksA in overall α-helical folding and a pair of conserved acidic residues at the flexible loop in the C-terminal coiled-coil domain^28^. In contrast to DksA proteins, which mostly control initiation of transcription^29^, Gre factors regulate the elongation step by coordinating a Mg^2+^ cation in the active site, which catalyzes water-mediated endonuclease cleavage of nascent transcripts backtracked into the secondary channel^30^. Because of the essential role DksA plays in the antioxidant defenses of *Salmonella*^6^ and because of the structural similarities between DksA and Gre factors, we have tested whether binding of Gre factors to RNA polymerase activates antioxidant programs that protect *Salmonella* against the antimicrobial activity engendered as NOX2 enzymatic complexes assemble at phagosomal membranes. Our investigations herein demonstrate that Gre factors control transcription elongation of EMP glycolysis and ETC genes, thereby balancing biosynthetic, energetic and redox outputs associated with increase *Salmonella* fitness during exposure to ROS in the innate host response.

## RESULTS

### Gre factors defend *Salmonella* against oxidative products of the phagocyte NADPH oxidase

The *Salmonella* genome encodes two homologous Gre proteins. Compared to wildtype controls, *Salmonella* mutants bearing deletions in *greA* or *greB* genes were slightly attenuated in an acute C57BL/6 murine model of infection (Fig. 1A). However, isogenic strains carrying deletions in both *greA* and *greB* genes (Δ*greAB*) were highly attenuated in C57BL/6 mice (Fig. 1A). Virulence of Δ*greAB Salmonella* could be complemented with either *greA* or *greB* genes expressed from the low copy plasmid pWSK29, demonstrating that both Gre factors play indispensable but mostly overlapping functions in *Salmonella* pathogenesis. *Salmonella* bearing deletions in both *greA* and *greB* genes remained attenuated in *nos2*^-/-^ mice lacking the inducible nitric oxide synthase, but recovered virulence in *nox2*^-/-^ mice deficient in the gp91*phox* subunit of the phagocyte NADPH oxidase (Fig. 1B). Δ*greA* and Δ*greB Salmonella* were hypersusceptible to H_2_O_2_ killing *in vitro* (Fig. 1C). Simultaneous elimination of *greA* and *greB* genes further sensitized *Salmonella* to the bactericidal (Fig. 1C) and bacteriostatic (Fig. 1D) activities of H_2_O_2_. Together, this research suggests that *Salmonella* have leveraged the regulatory activity of Gre factors to resist oxidative stress in the innate host response.

**Fig. 1.**
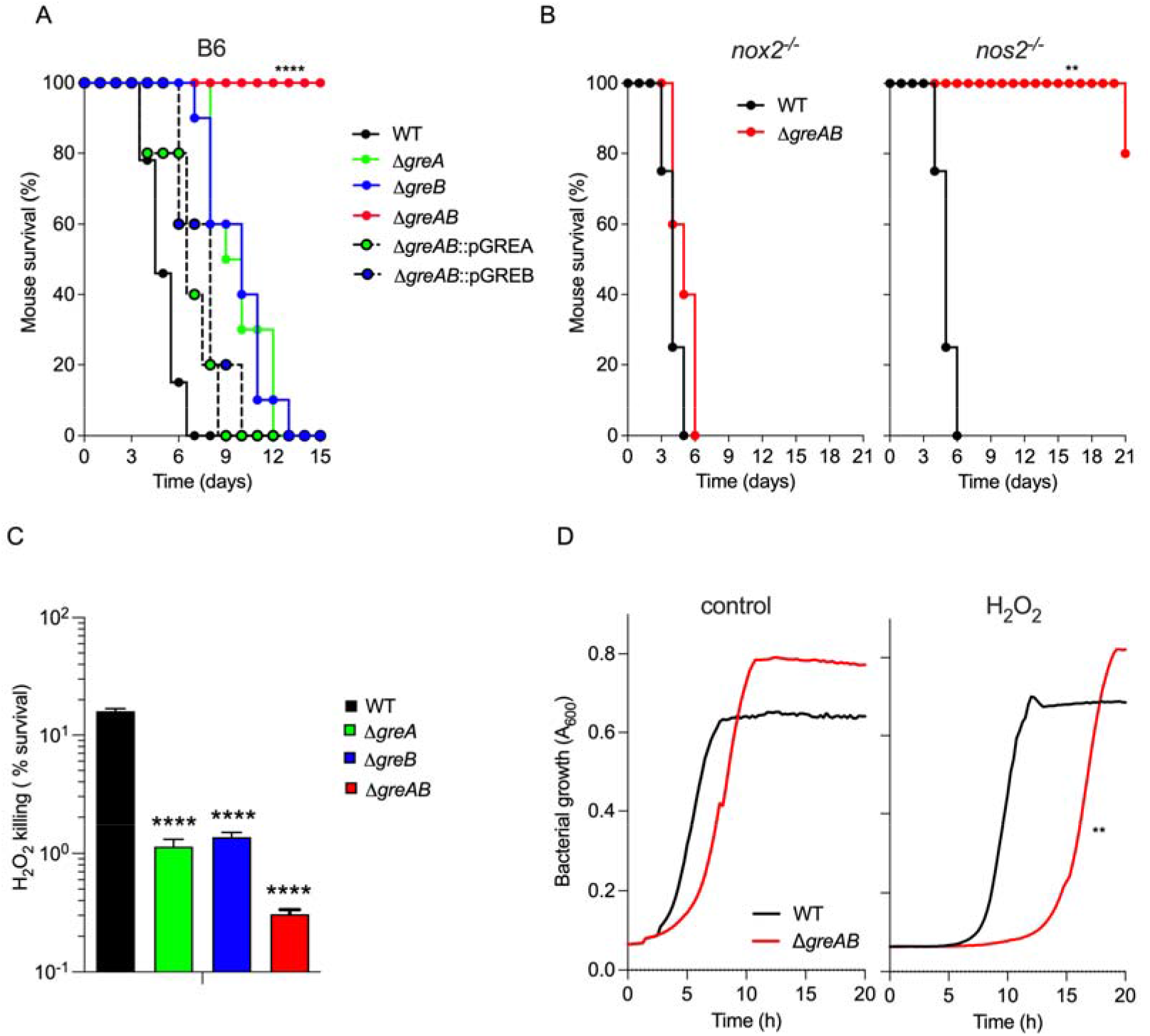
Gre factors regulate resistance of *Salmonella* to the antimicrobial activity of NOX2. Survival of C57BL/6 (A), *nox2^-/-^* and *nos2^-/-^* (B) mice after i.p. inoculation of 100 CFU of the indicated strains of *Salmonella* (N = 10). Statistical differences (*p*<0.0001) were calculated by logrank analysis. The Δ*greAB* mutant was complemented with either *greA* or *greB* genes expressed from their native promoters in the low copy number pWSK29 plasmid (i.e., pGREA and pGREB, respectively). (C) Killing of *Salmonella* after 2h of treatment with 400 μM H_2_O_2_. Bacterial cultures were grown overnight in LB broth, diluted to 2×10^5^ CFU/ml in PBS and treated for 2h with 400 μM H_2_O_2_. Killing, expressed as percent survival compared to the bacterial burden at time zero, is the mean ± S.D. (N=12-16); *p*<0.0001 as determined by one-way ANOVA. (D) Growth of wild-type (WT) and Δ*greAB* mutant *Salmonella* in EG minimal medium in the presence or absence of 200 μM H_2_O_2_ as measured in a Bioscreen. Data are shown as the mean (N=4-5). \*\**p*<0.01 as determined by *t*-test.

### *Salmonella* require Gre factors to grow on glucose

We noticed that Δ*greAB Salmonella* grow poorly in glucose (Fig. 1D, 2A). Because glycolysis plays a salient role in resistance of *Salmonella* to the antimicrobial activity of the phagocyte NADPH oxidase^5^, we sought to examine the underlying causes that prevent Δ*greAB Salmonella* from effectively utilizing glucose. Interestingly, an extended lag phase was also observed when Δ*greAB Salmonella* were grown in MOPS minimal medium supplemented with either fructose, pyruvate, succinate, fumarate or malate (Fig. S1A-J), suggesting that Gre factors regulate assimilation of a variety of sugars. The defective growth of Δ*greAB Salmonella* in MOPS-glucose (GLC) medium was ameliorated under anaerobic conditions (Fig. 2A). Likewise, supplementation of MOPS-GLC medium with GSH partially restored the growth of aerobic Δ*greAB Salmonella* to wildtype levels (Fig. 2B). Wildtype and Δ*greAB Salmonella* grew with similar kinetics in MOPS minimal medium supplemented with either Casamino acids or an equal mix of all 20 amino acids (Fig. 2C), suggesting that the deletion of Gre factors may induce functional amino acid auxotrophies. To test this idea further, we quantified amino acid pools in wildtype and Δ*greAB Salmonella* (Fig. 2D, S2). Most amino acids were at similar or even higher concentrations in Δ*greAB Salmonella* compared to wildtype bacteria. However, compared to wildtype controls, Δ*greAB Salmonella* harbored lower intracytoplasmic concentrations of the aromatic amino acids phenylalanine and tyrosine, which are generated from erythrose 4-phosphate and phosphoenolpyruvate, and valine, which is synthesized from pyruvate (Fig. 2D). These findings indicate that *Salmonella* lacking Gre factors suffer from amino acid bradytrophies. This idea is supported by the observation that Δ*greAB Salmonella* contained higher levels of the nucleotide alarmone ppGpp (Fig. 2E), which can be produced by RelA proteins sensing an upsurge of deacylated tRNAs^31^.

**Fig. 2.**
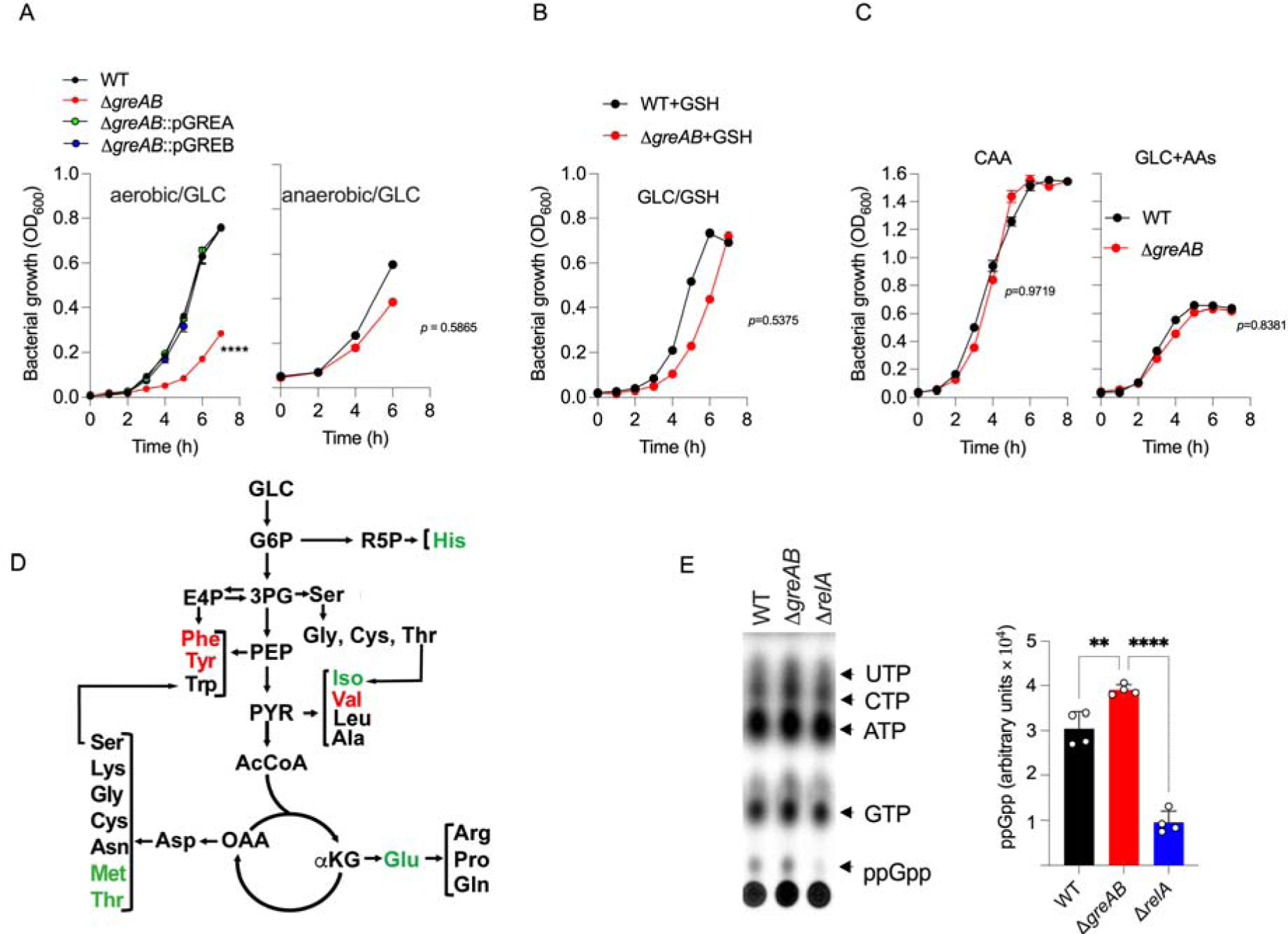
Gre factors facilitate growth of *Salmonella* in glucose. (A) Aerobic and anaerobic growth of wildtype (WT) and Δ*greAB* mutant *Salmonella* in MOPS-GLC medium. Where indicated, the Δ*greAB* mutant was complemented with the *greA* or *greB* genes. *p*<0.0001 as determined by two-way ANOVA. (B) Aerobic growth of *Salmonella* strains in MOPS-GLC medium supplemented with 2 mM GSH (B), or MOPS minimal medium supplemented with 0.4% casamino acids (CAA) or GLC containing 40 μg/ml each of the 20 amino acids (AAs) (C). Data in A-C are mean ± S.D. from at least three biological replicates. (D) Schematic representation of amino acid biosynthesis from glycolysis and TCA cycle intermediates. *Salmonella* strains grown in MOPS-GLC medium to an OD_600_ of 0.25 were collected for amino acid quantification by liquid chromatography mass spectrometry (LC-MS). Amino acids that were statistically (*p*<0.01) more and less abundant in Δ*greAB* than WT controls are shown in green and red, respectively. AAs contained at similar levels in Δ*greAB* and WT *Salmonella* are shown in black. Data are the mean ± S.D. (N=5). (E) Analysis for nucleotide biosynthesis. TLC autoradiogram (Left panel) and densitometry (right panel) of ^32^P-labeled nucleotides extracted from an OD_600_ of 0.25 of *Salmonella* strains grown in MOPS-GLC media. Data are shown as mean ± S.D. (N=4). **,**** *p*< 0.01 and *p*< 0.0001, respectively, as determined by one-way ANOVA. The online version of this article includes the following figure supplements and source data for figure 4: Figure Supplement S1. Effect of carbon source on *Salmonella* growth. Figure Supplement S2. Amino acid pools in *Salmonella* grown on glucose. Source data: GEO#GSE197076

### Gre factors activate transcription of glycolysis and electron transport chain genes

To search into the mechanisms underlying the poor growth of Δ*greAB Salmonella* on glucose as sole carbon source, we compared the transcriptional profiles of log phase wildtype and Δ*greAB Salmonella* grown in MOPS-GLC medium (Fig. 3A). Principal component analysis indicated that wildtype and Δ*greAB Salmonella* grown on glucose markedly differ in their transcriptome (Fig. S3)(GEO#GSE203342). The Δ*greAB Salmonella* strain showed defective expression of *gapA*, *eno*, and *pykF* glycolytic genes encoding glyceraldehyde dehydrogenase (GAPDH), enolase and pyruvate kinase, respectively (Fig.3A). RT-PCR analysis validated these findings (Fig. S4). Δ*greAB Salmonella* grown in MOPS-GLC medium harbored less (*p* < 0.05) GAPDH enzymatic activity than wildtype controls (Fig. 3B). The Δ*greAB Salmonella* strain seems to resolve the reduced carbon flow through lower glycolysis by upregulating transcription of the *talA*-encoded aldolase that shuttles carbon from upper glycolysis into the pentose phosphate pathway (Fig. 3A, S4). Moreover, Δ*greAB Salmonella* overexpressed the Entner-Doudoroff gene *dgaF* encoding 2-dehydro-3-deoxy-phosphogluconate aldolase (Fig. 3A). The Δ*greAB Salmonella* strain supported normal or increased expression of oxidative TCA genes involved in α-ketoglutarate synthesis, which may explain the accumulation of abnormally high concentrations of glutamic acid, citrulline and ornithine in Δ*greAB Salmonella* grown on glucose (Fig. 2D, S2).

**Fig. 3.**
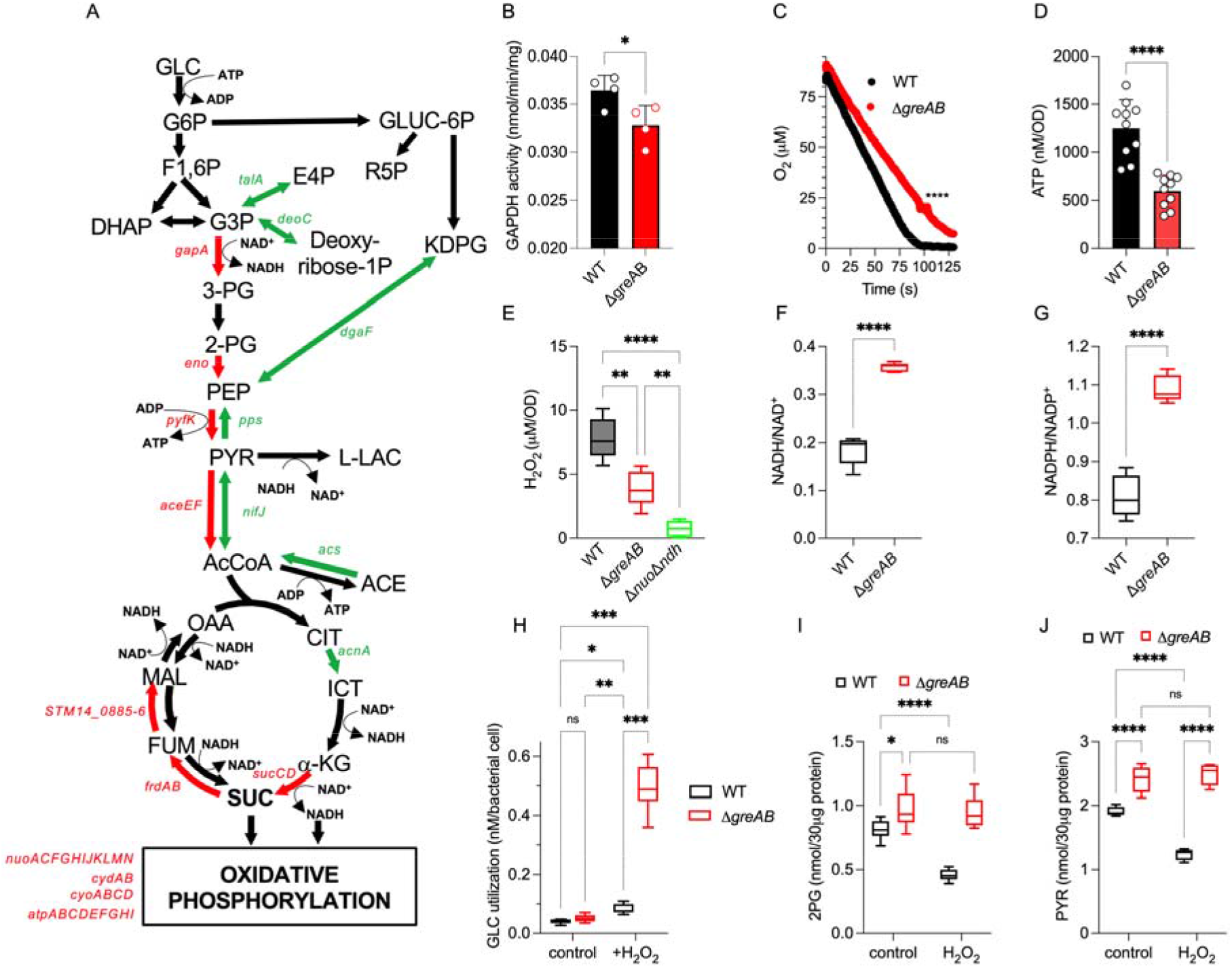
Gre factors activate transcription of EMP glycolysis and ETC genes. (A) Schematic representation of transcription of glycolytic, TCA and oxidative phosphorylation genes in WT and Δ*greAB* mutant *Salmonella*. Red and green arrows show significantly down and up-regulated gene transcripts, respectively in Δ*greAB Salmonella* compared to WT controls, as determined by RNA-Seq of *Salmonella* grown in MOPS-GLC medium to an OD_600_ of 0.25. Highlighted genes undergo at least one two-fold change and have *q* < 0.05 (N=4) (B) GAPDH activity in WT and Δ*greAB* mutant *Salmonella* grown in MOPS-GLC medium to OD_600_ of 0.25. Data are the mean ± S.D (N=4). Consumption of O_2_ (C) and endogenous H_2_O_2_ synthesis (E) by exponentially grown *Salmonella* were measured polarographycally in an ISO-OXY/HPO analyzer equipped with an O_2_ or H_2_O_2_ probes, respectively. Data are the mean ± SD (N=5-6). Intracellular ATP concentrations (D), nicotinamide adenine nucleotide ratios (F, G), glucose (GLC) consumption (H), as well as 2-phosphoglycerate (2PG) (I) and pyruvate (PYR) (J) concentrations were recorded in *Salmonella* grown to an OD_600_ of 0.25 in MOPS-GLC media. Where indicated, samples were treated with 400 μM H_2_O_2_. Data in D are shown as mean ± S.D. (N=10), F and G (N=10), H-I (N=6-8). *, **, ***,**** *p* < 0.05, *p*< 0.01, *p*< 0.001 and *p*< 0.0001, respectively as determined by *t*-test (B, C, D, F and G), one-way ANOVA (E), or two-way ANOVA (H-J). The online version of this article includes the following figure supplements for figure 3: Figure Supplement S3. RNA seq analysis of *Salmonella* grown in glucose. Figure Supplement S4. Gene expression in *Salmonella* grown on glucose. Figure Supplement S5. H_2_O_2_ consumption by *Salmonella* and transcriptional pausing.

The absence of Gre factors also resulted in the downregulation of genes within the ETC, including *nuo*, *cyd*, *cyo*, and *atp* operons encoding NDH-I NADH dehydrogenase, high and low affinity quinol oxidases, and ATP synthase, respectively (Fig. 3A, S4). In agreement with the transcriptional profiles, Δ*greAB Salmonella* sustained lower aerobic respiration (Fig. 3C) and harbored diminished ATP pools (Fig. 3D) compared to wildtype bacteria.

Thus far, our data indicate that *Salmonella* lacking Gre factors are hypersusceptible to oxidative stress, sustain decrease glycolytic activity, and exhibit poor aerobic respiration. Because the ETC is a notorious source of endogenous oxidative stress, we sought to compare the adventitious production of ROS in wildtype and Δ*greAB Salmonella*. A Δ*greAB Salmonella* strain accumulated lower concentrations of H_2_O_2_ than wildtype *Salmonella* (Fig. 3E). A Δ*nuo* Δ*ndh* strain lacking both NADH dehydrogenases engendered trace amounts of H_2_O_2_, suggesting that NADH dehydrogenases are the main source of endogenous oxidative stress in *Salmonella*. Moreover, Δ*greAB Salmonella* expressed significantly (*p* < 0.0001) more catalase activity than wildtype bacteria (Fig. S5A). We were also surprised by the excellent reductive power of Δ*greAB Salmonella*. First, Δ*greAB Salmonella* harbored a significantly (*p* < 0.0001) higher NADH/NAD^+^ ratio than wildtype controls (Fig. 3F), likely reflecting reduced transcription of NADH dehydrogenases and aerobic respiration genes. Second, Δ*greAB* mutant harbored a significantly (*p* < 0.0001) higher NADPH/NADP^+^ ratio than wildtype controls (Fig. 3G), consistent with the transcriptional analysis that showed overutilization of the pentose phosphate pathway (Fig. 3A). Importantly, NADPH fuels classical antioxidant defenses such as GSH and thioredoxin. Treatment of *Salmonella* with H_2_O_2_ increased glucose utilization in both wildtype and Δ*greAB Salmonella*, and the Δ*greAB* mutant consumed higher glucose than wildtype controls after treatment with H_2_O_2_ (Fig. 3H). Wildtype *Salmonella* accumulated lower concentrations of 2-phosphoglycerate (Fig. 3I) and pyruvate (Fig. 3J) upon H_2_O_2_ treatment. Δ*greAB Salmonella* contained higher levels of 2-phosphoglycerate and pyruvate than wildtype controls, and treatment of this mutant with H_2_O_2_ did not alter the intracellular concentrations these glycolytic intermediates.

Collectively, these investigations indicate that the transcriptional control Gre factors exert on glycolytic and ETC genes aids *Salmonella* to allocate carbon, and maintain energy and redox balance across substrate-level and oxidative phosphorylation. Furthermore, these findings suggest that the hypersusceptibility of Δ*greAB Salmonella* to oxidative stress must be independent of their ability to handle H_2_O_2_ by classical detoxification systems.

### Gre factors relieve transcriptional pauses of glycolytic and aerobic respiration genes

We next examined the mechanism underlying the control of glycolytic and aerobic gene transcription by Gre factors. The addition of GreA or GreB recombinant proteins (Fig. S5B) to a reconstituted *in vitro* system increased expression of *gapA* (Fig. 4A) and *cydA* (Fig. 4B) genes. These data demonstrate that Gre factors directly activate transcription of both *gapA*, the first gene in the payoff phase of glycolysis, and the *cydAB* operon encoding cytochrome *bd,* which imparts resistance to H_2_O_2_, NO and hydrogen sulfide^32–34^. To further probe the mechanism by which Gre factors promote *gapA* transcription, we visualized the products of the *in vitro* transcription assays on urea PAGE gels. RNA polymerase paused at several sites between the +15 and +26 positions from the *gapA* transcription start site (Fig. 4C). Inclusion of Gre factors, especially GreB, in the *in vitro* transcription reactions resolved the transcriptional pauses in the *gapA* gene. Gre factors also resolved transcriptional pauses in the EMP *eno* gene and the aerobic *cydA* gene (Fig. 4D, 4E). DksA, which also binds to the secondary channel of RNA polymerase, did not resolve transcriptional pausing in the *eno* gene (Fig. S5C). Both GreA and GreB proteins increased both *gapA* and *cydA* transcriptional runoff products (Fig. 4C, 4E). Together, these investigations identify the Gre-dependent rescue of transcriptional pausing as a vital regulatory step in the activation of transcription of glycolytic and respiratory genes in *Salmonella*.

**Fig. 4.**
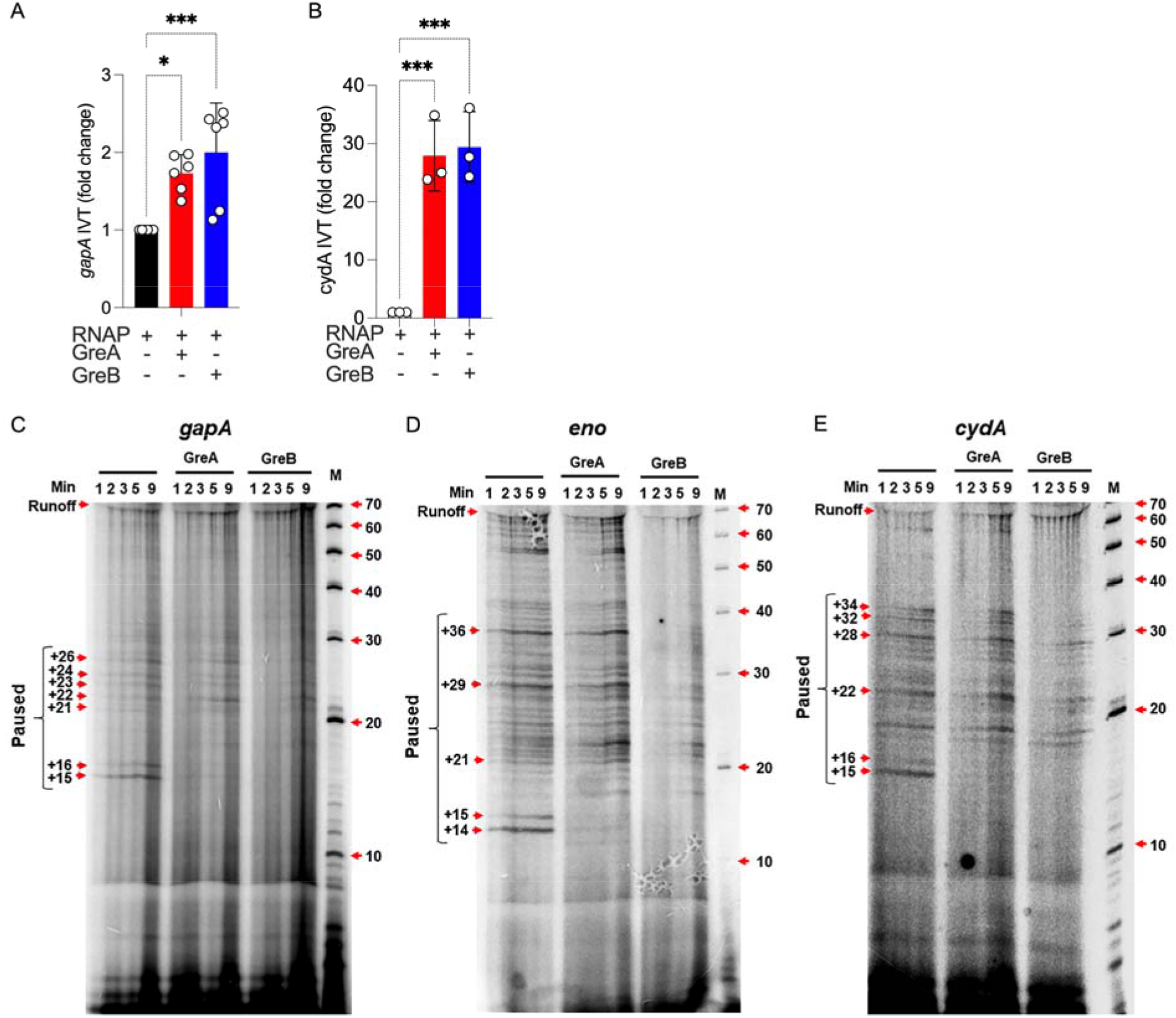
Direct regulation of gene transcription by Gre factors. Effect of the addition of Gre proteins in the *in vitro* transcription of *gapA* (A) and *cydA* (B) genes in a reconstituted biochemical system. Data are shown as mean ± S.D. (N=6 for *gapA* and 3 for *cydA*). *, *** *p* < 0.05 and *p*< 0.001, respectively as determined by one-way ANOVA. (C-E) Pausing of *in vitro* transcription reactions containing *gapA*, *eno* or *cydA* templates was visualized in urea PAGE gels of RNA products labeled with α^32^P-UTP. Representative blots from 3 independent experiments. The online version of this article includes the following figure supplements for figure 4: Figure Supplement S5. H_2_O_2_ consumption by *Salmonella* and transcriptional pausing.

### The ETC allows *Salmonella* to grow on glucose and withstand oxidative stress

Our investigations have shown that Gre factors activate the transcription of EMP glycolysis and ETC genes. Glycolytic defects may contribute to the susceptibility of Δ*greAB* mutant *Salmonella* to oxidative killing, as suggested by the hypersusceptibility of Δ*gpmA Salmonella* lacking phosphoglycerate mutase to H_2_O_2_ and NOX2-dependent killing^5^. In the following investigations, we tested whether aerobic respiration may also be a critical component by which *Salmonella* grow on glucose and resist NOX2-mediated host defense. *Salmonella* strains bearing deletions in either *cydAB* or *atpD* genes encoding subunits of cytochrome *bd* and ATP synthase grew poorly in MOPS-GLC medium, and a Δ*nuo* Δ*ndh* mutant lacking NDH-I NADH dehydrogenase completely failed to grow on glucose (Fig. 5A). These findings demonstrate that the ETC and oxidative phosphorylation boost the glycolytic capacity of *Salmonella*, and suggest that decreased expression of ETC genes may contribute to the glycolytic defects of Δ*greAB Salmonella*.

**Fig. 5.**
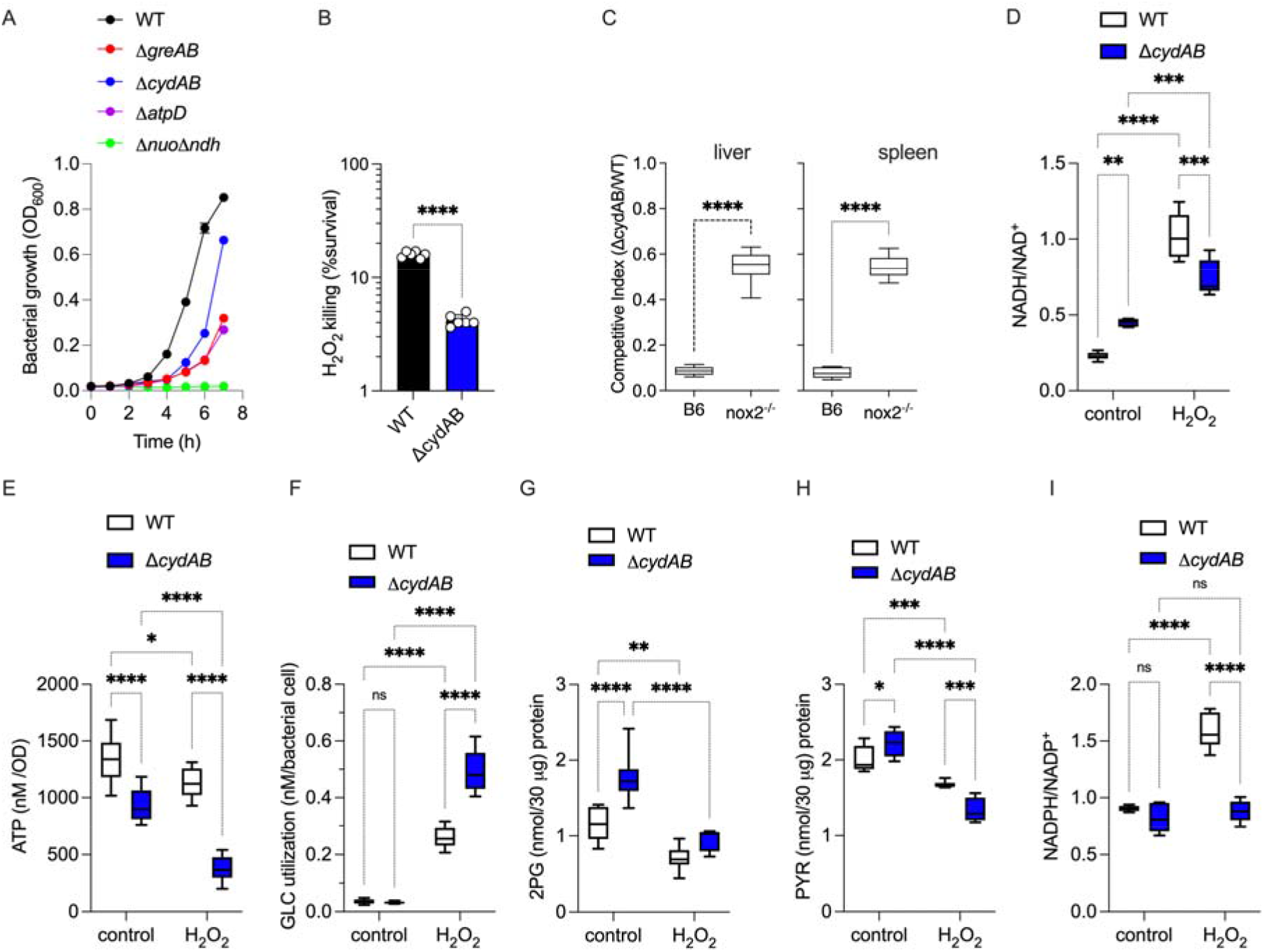
Contribution of aerobic respiration to the antioxidant defenses of *Salmonella*. (A) Aerobic growth of *Salmonella* strains in MOPS-GLC media. Data are the mean ± S.D (N=3) (B) H_2_O_2_ killing of *Salmonella*. Bacterial cultures were grown overnight in LB broth, diluted to 2×10^5^ CFU/ml in PBS and treated for 2h with 400 μM H_2_O_2_. Killing is expressed as percent survival compared to the bacterial burden at time zero. N=6; *p*<0.0001 as determined by unpaired *t*-test. (C) Competitive index of *Salmonella* in livers and spleen of C57BL/6 and *nox2*^-/-^ mice after i.p. inoculation with 100 CFU of equal numbers of WT and Δ*cydAB Salmonella* (n = 10). Statistical differences (****, *p*<0.0001) were calculated by unpaired *t*-test. Intracellular nicotinamide adenine nucleotide ratios (D, I), ATP concentrations (E), glucose utilization (F), 2-PG (G) and pyruvate (H) production were estimated in *Salmonella* grown to an OD_600_ of 0.25 in MOPS-GLC media. Selected samples were treated with 400 μM H_2_O_2_ for 2h for glucose utilization data and 30 minutes for others. Data are the mean ± S.D from at least six to ten biological replicates. *, **, ***, ****, *p*< 0.05, *p*< 0.01, *p*< 0.001 and *p*< 0.0001, respectively as determined by two-way ANOVA.

To test the importance of aerobic respiration in the resistance of *Salmonella* to oxidative stress, we used a Δ*cydAB* mutant lacking the quinol oxidase cytochrome *bd*. A *Salmonella* strain bearing mutations in the *cydAB* operon was hypersensitive to H_2_O_2_ killing (Fig. 5B), and was attenuated in immunocompetent C57BL/6 mice (Fig. 5C). The attenuation of Δ*cydAB Salmonella* was partially reversed in *nox2*^-/-^ mice (Fig. 5C), demonstrating that aerobic respiration is a previously unappreciated aspect of the antioxidant defenses of *Salmonella*. Cytochrome *bd* has peroxidatic activity^32^, raising the possibility that deletion of the *cydAB* operon could predispose *Salmonella* to oxidative stress by the corresponding diminished capacity to detoxify H_2_O_2_. However, wildtype and Δ*cydAB Salmonella* detoxified H_2_O_2_ with apparently similar (*p* > 0.05) kinetics (Fig. S5D). Therefore, we compared several metabolic signatures associated with resistance to oxidative stress between wildtype and Δ*cydAB Salmonella*. Wildtype *Salmonella* contained a lower NADH/NAD^+^ ratio than Δ*cydAB* controls (Fig. 5D), consistent with the greater capacity of the former to perform aerobic respiration. The elevated NADH/NAD^+^ ratios recorded in Δ*cydAB Salmonella* may explain the poor growth of this strain in glucose (Fig. 5A), as NAD^+^ is a cofactor of GAPDH in EMP glycolysis. H_2_O_2_ treatment elevated the NADH/NAD^+^ ratio in both wildtype and Δ*cydAB Salmonella* (Fig. 5D), probably as a consequence of the oxidative damage of NDH-I NADH dehydrogenase^5^ and the downregulation of the *nuo* operon (Fig. 3A, S4). Together, these investigations suggest that regeneration of NAD^+^ needed for glycolytic activity cannot completely be satisfied in lactate and succinate fermentation^5^, but requires the oxidizing activity of NADH dehydrogenases in the ETC.

The Δ*cydAB* strain harbored reduced ATP content compared to wildtype controls, and both strains suffered reductions in ATP upon H_2_O_2_ treatment (Fig. 5E). At the culture density used in these experiments, 400 μM H_2_O_2_ was similarly bacteriostatic for both wildtype and Δ*cydAB Salmonella* (Fig. S5E). H_2_O_2_ increased glucose utilization in both wildtype and Δ*cydAB Salmonella*, but diminished production of 2-phosphoglycerate and pyruvate (Fig. 5F-H). The decreased carbon flow through lower glycolysis may stem from the oxidation of the catalytic cysteine in GAPDH, favoring usage of the pentose phosphate pathway. In support of this idea, the NADPH/NADP^+^ ratio was significantly (*p* < 0.0001) increased in wildtype *Salmonella* upon treatment with 400 μM H_2_O_2_ (Fig. 5I). Interestingly, Δ*cydAB Salmonella* did not experience similar increases in NADPH/NADP^+^ after the addition of H_2_O_2_. Together, these investigations suggest that aerobic respiration allows for optimal utilization of glycolysis and the pentose phosphate pathway, thus contributing to the antioxidant defenses that protect *Salmonella* against NOX2-mediated host immunity. Consequently, the lower aerobic respiration recorded in Δ*greAB Salmonella* may contribute to the hypersusceptibility of this strain to ROS.

### *Salmonella* undergoing oxidative stress utilize the methylglyoxal pathway

H_2_O_2_ diminished redox potential in both wildtype and Δ*greAB Salmonella*, as indicated by increased ratiometric roGFP measurements (Fig. 6A). This approach also showed that the cytoplasm of untreated Δ*greAB Salmonella* is heavily oxidized compared to wildtype controls. We were surprised by these puzzling findings because Δ*greAB Salmonella* synthesize lower H_2_O_2_ and contain higher ratios of reduced nicotinamide adenine dinucleotides than wildtype *Salmonella* (Fig. 3E, 3F). RoGFP fluorescence reflects the GSH/GSSG redox buffering of the cell^35^. In agreement with the roGFP estimations, Δ*greAB Salmonella* contained a significantly (*p* < 0.0001) lower concentration of reduced GSH than wildtype controls (Fig. 6B). Loss of GSH in Δ*greAB Salmonella* was not paralleled by gains in oxidized GSH (Fig. 6B). Based on these seemingly paradoxical observations, we turned our attention to the methylglyoxal pathway, which branches off from glycolysis and generates the aldehyde methylglyoxal during the recycling of glycolytic phosphosugar intermediates and consumes GSH (Fig. 6C). Δ*greAB Salmonella* harbored abnormally high concentrations of aldehyde (Fig. 6D) and D-lactate (Fig. 6E). These findings indicate that Δ*greAB Salmonella* is wired to overutilize the methylglyoxal pathway, providing a reasonable explanation for the fact that this mutant has low concentrations of GSH but apparently wildtype levels of oxidized GSH. Treatment of wildtype *Salmonella* with H_2_O_2_ increased production of aldehyde and D-lactate (Fig. 6D, 6E). These findings indicate that *Salmonella* activate the methylglyoxal pathway following H_2_O_2_ treatment. Recycling of GSH from the S-lactoyl-glutathione intermediate by the *gloB*-encoded glyoxylase was critical for growth of *Salmonella* on glucose (Fig. 6F). Importantly, a Δ*gloB* mutant was attenuated in C57BL/6 mice, but regained virulence in *nox2*^-/-^ mice lacking NADPH oxidase (Fig. 6G), demonstrating that the formation of GSH through the cleavage of S-lactoyl-glutathione contributes to the antioxidant defenses of *Salmonella*.

**Fig. 6.**
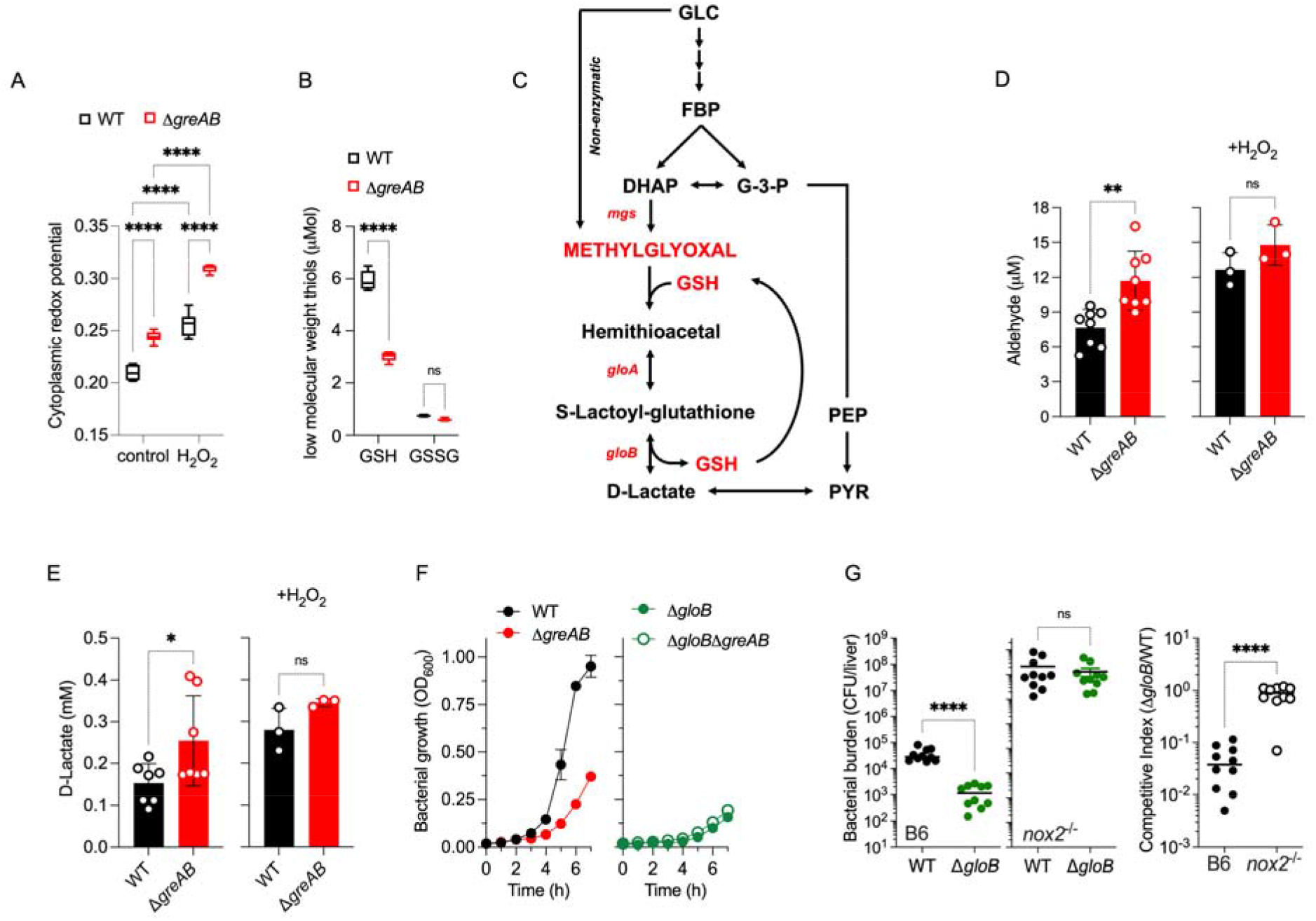
Oxidative stress stimulates carbon flux through the methylglyoxal pathway in *Salmonella.* (A) Cytoplasmic redox potential was estimated by recording the fluorescent spectra of roGFP2 in bacteria grown in MOPS-GLC medium to OD_600_ of 0.25. Some of the bacterial cultures were treated for 1 min with 400 μM H_2_O_2_. The relative redox potential denotes the ratio of roGFP2_ox_ /roGFP2_red_ emission signals at 510 nm after excitation at 405 and 480 nm. (B) Low-molecular weight thiols (LMWT) were estimated by liquid chromatography mass spectrometry (LC-MS). N=5-6. ****, *p*< 0.0001 as assessed by two-way ANOVA. (C) Schematic representation of methylglyoxal pathway in *Salmonella.* Intracellular (D) aldehyde and (E) D-lactate in *Salmonella* grown to an OD_600_ of 0.25 in MOPS-GLC media. Selected samples were treated with 400 μM H_2_O_2_. Data are the mean ± S.D. (N=3-8). *, **, *p*< 0.05 and *p*< 0.01, respectively as determined by unpaired *t*-test. (F) Growth of the indicated *Salmonella* strains in MOPS-GLC media. Data are the mean ± S.D (N=3). (G) Bacterial burden and competitive index of *Salmonella* in livers of C57BL/6 and *nox2*^-/-^ mice after i.p. inoculation with 100 CFU of equal numbers of WT and Δ*gloB Salmonella* (N = 10). Statistical differences (****, *p*<0.0001) were calculated by unpaired *t*-test.

## DISCUSSION

Transcriptional pauses that arise from the misincorporation of nucleoside monophosphates into the growing elongation complex are resolved by the Gre-directed endonuclease activity of RNA polymerase^30, 36, 37^. In addition to preserving transcriptional fidelity, reactivation of stalled ternary elongation complexes by Gre factors reduces conflicts with the replisome^38–40^, likely supporting bacterial growth^40, 41^. Our investigations provide a non-mutually exclusive alternative for the growth-promoting activity of Gre factors. Gre-mediated transcriptional activation of EMP glycolysis and ETC accomplishes the biosynthetic, energetic and redox demands of the cell. The underutilization of both glycolysis and aerobic respiration in Δ*greAB Salmonella* results in amino acid bradytrophies, shortages in energy supply, as well as the puzzling coexistence of reductive and electrophilic stress. The adaptive utilization of pentose phosphate and Entner-Doudoroff pathways in Δ*greAB Salmonella* incompletely fulfills energetic and biosynthetic demands. The Gre-dependent transcriptional activation of EMP glycolysis and ETC helps *Salmonella* effectively utilize glucose, thereby meeting the metabolic needs of *Salmonella* in the host.

Our investigations demonstrate that the equilibrated apportioning of resources to EMP and ETC that follows the transcriptional regulation afforded by Gre factors helps *Salmonella* resist oxidative stress. We were rather surprised by the high NADH/NAD^+^ and NADPH/NADP^+^ reducing power, as well as by the excellent catalase enzymatic activity recorded in Δ*greAB Salmonella*. Moreover, the absence of Gre factors decreases aerobic respiration, thereby diminishing endogenous production of ROS. Despite these excellent predictors of bacterial resistance to oxidative stress, Δ*greAB Salmonella* lack the tolerance of wildtype cells to the bacteriostatic and bactericidal activity of ROS and are attenuated in NOX2 proficient mice. This research is consistent with the idea that glycolysis is a central aspect of the adaptive response of *Salmonella* to the antimicrobial activity of the phagocyte NADPH oxidase^5^. Our investigations have also revealed the unexpected finding that, although diminished during periods of oxidative stress, aerobic respiration is still critical for both growth on glucose and resistance to oxidative killing. Thus, resistance to oxidative stress cannot be met with overflow metabolism alone, but requires the additional redox balancing and energetic outputs associated with aerobic respiration. We conclude that the metabolic adaptations that follow the resolution of transcriptional pausing at EMP and ETC genes is a *sine qua non* for resistance of *Salmonella* to oxidative stress, and that classical antioxidant defenses cannot overcome the susceptibility of bacteria bearing inappropriate metabolic signatures to ROS.

Rapidly growing cells turn on metabolic overflow, an adaptive response that favors deployment of glycolysis and fermentation in place of oxidative phosphorylation^42, 43^. Two models have been put forward to explain the advantages associated with overflow metabolism during exponential grow. First, fast growing cells allocate proteome resources to less demanding glycolysis, fermentation and the Entner-Doudoroff pathway, while simultaneously reducing proteomic allotment to the resource-intensive ETC^42^. As glycolysis generates a lower energic output than aerobic respiration, the cell must compensate by increasing carbon uptake and breakdown. Second, the growing cell suffers from a “membrane real estate” crisis^44, 45^. As the surface to volume ratio diminishes in the growing cell, the availability of a lipid scaffold becomes limiting for the assembly of ETC protein complexes. In analogy to rapidly growing bacteria, increases in glucose utilization and NADH/NAD+ ratios and decreases in ATP noted in H_2_O_2_-treated *Salmonella* indicate that oxidative stress induces overflow metabolism. Following exposure to ROS, *Salmonella* diminish oxidative phosphorylation while upregulating substrate-level phosphorylation (this paper and ref^5^). The induction of overflow metabolism in bacterial cells undergoing oxidative stress seems to be unrelated to the membrane real state crisis of fast-growing cells, because *Salmonella* exposed to sublethal concentrations of H_2_O_2_ discontinue growth. Instead, oxidative stress seems to trigger a quinone crisis. Shifting some of the energetic and redox production away from the respiratory chain during oxidative stress allows the Dsb system to deliver electrons to oxidized quinones, thus powering the isomerization of erroneously generated disulfide bonds in periplasmic proteins by exogenous ROS^5^. The NADH dehydrogenase encoded in the *nuo* operon acts as a ROS sensor that switches on overflow metabolism in *Salmonella* experiencing oxidative stress^5^.

The downregulation of oxidative phosphorylation creates an energetic and redox balancing dilemma in bacteria undergoing oxidative stress. *Salmonella* resolve this conundrum by increasing glucose utilization, which generates ATP in both lower glycolysis and the fermentation of pyruvate to acetate. Increased glycolytic activity also boosts NADPH synthesis in pentose phosphate pathway that fuels classical antioxidant defenses, and supplements the shrinking redox balancing capacity of an oxidatively damaged ETC by redirecting carbon to lactate fermentation and the reductive branch of TCA^5^. Overflow metabolism, however, generates a poisonous intermediate. To maintain energetic and redox outputs with less productive glycolysis and fermentation, bacteria are forced to liberate inorganic phosphate from phosphosugar intermediates arising as glycolysis is cranked up. Our data show that *Salmonella* solve this predicament by upregulating the methylglyoxal pathway. Detoxification of phosphosugars generates the electrophilic intermediate methylglyoxal, an aldehyde responsible for the formation of advance glycation end-products^46^. Methylglyoxal is detoxified by the glyoxylase system in a reaction that consumes GSH. Ironically, GSH is not only consumed by the nucleophilic avidity of H_2_O_2_ for the tripeptide, but more importantly by the electrophilic attack of methylglyoxal produced as ROS-stressed cells crank up glycolysis. Despite the potential cytotoxicity of this metabolic pathway, the role the methylglyoxal pathway plays in glucose utilization offers *Salmonella* an adaptive advantage against the oxidative stress generated by NOX2 in the innate host response.

Most studies have focused on the regulation of metabolism that follows the hierarchical control provided by transcriptional activators as well as the allosteric regulation of central metabolic enzymes by metabolites and posttranslational modifications. In complement to these studies, our investigations demonstrate that transcription elongation is key for balanced metabolic outputs. Gre-dependent transcriptional elongation of EMP and ETC genes balances the simultaneous usage of overflow metabolism and aerobic respiration, thus fulfilling the biosynthetic, energetic and redox requirements that help *Salmonella* withstand the antimicrobial activity of NOX2 in the acute host response.

## MATERIALS AND METHODS

### Bacterial strains, plasmids and growth conditions

The *Escherichia coli* strains DH5α and BL21(DE3)pLysS were grown in Luria–Bertani (LB) broth or agar at 37°C. *Salmonella enterica* serovar *Typhimurium* strain 14028s (ATCC, Manassas, VA) and its mutant derivatives were grown in LB broth or E-salts minimal medium [57.4 mM K_2_HPO_4_, 1.7 mM MgSO_4_, 9.5 mM citric acid and 16.7 mM H_5_NNaPO_4_, pH 7.0, supplemented with 0.1% casamino acids, and 0.4% D-glucose (EGCA)], or MOPS minimal medium [40 mM MOPS buffer, 4 mM Tricine, 2 mM K_2_HPO_4_, 10 μM FeSO_4_·7H_2_O, 9.5 mM NH_4_Cl, 276 μM K_2_SO_4_, 500 nM CaCl_2_, 50 mM NaCl, 525 μM MgCl_2_, 2.9 nM (NH_4_)_6_Mo7O_24_·4H_2_O, 400 nM H_3_BO_3_, 30 nM CoCl_2_, 9.6 nM CuSO_4_, 80.8 nM MnCl_2_, and 9.74 nM ZnSO_4_, pH 7.2] supplemented with 0.4% D-glucose or 0.4% Casamino acids at 37°C in a shaking incubator. Ampicillin (100 μg/ml), kanamycin (50 μg/ml), chloramphenicol (20 μg/mL) and tetracycline (20 μg/mL) were used where appropriate.

### Construction of *Salmonella* Δ*greAB* mutants and complementation

Deletion mutants were constructed using the λ-Red homologous recombination system^47^. Specifically, the chloramphenicol cassette from the pKD3 plasmid and kanamycin cassette from pKD13 were PCR amplified using primers with a 5’-end overhang homologous to the bases following the ATG start site and the bases preceding the stop codon of *greA*, *greB, msgA, gloA* and *gloB* genes, respectively (Table S1). The PCR products were gel purified, and electroporated into *Salmonella* expressing the λ Red recombinase from the plasmid pTP233. Transformants were selected on LB plates containing 10 μg/ml chloramphenicol or 50 μg/ml kanamycin. To construct a mutant deficient in both Gre factors, the **Δ***greB*-Km mutation was moved into the **Δ***greA-*Cm mutant via P22-mediated transduction, and the pseudolysogens were eliminated by streaking on Evans blue uridine agar plates. Transformants were selected on 50 μg/ml kanamycin and 20 μg/ml chloramphenicol LB agar plates. The mutants were confirmed by PCR and sequencing.

The **Δ***greAB* mutant was complemented with *greA* or *greB* genes expressed from the low-copy pWSK29 plasmid^48^. The *greA* and *greB* coding regions plus a 400 bp upstream region including the native promoter were PCR-amplified using *greA* pro F and *greA* R or *greB* pro F and *greB* R primers, respectively (Table S2). The amplified PCR products were directly cloned into the SacII and BamHI restriction sites at the MCS of the pWSK29 vector. The resulting **Δ***greAB::greA* or **Δ***greAB::greB* complemented strains were selected on 250 μg/ml penicillin LB agar plates.

### Cloning, expression and purification of proteins

Recombinant 6XHis-tag GreA or 6XHis-tag GreB were produced by cloning *greA* and *greB* genes into NdeI and BamHI sites of the pET14B vector (Novagen) using *greA* F and *greA* R or *greB* F and *greB* R primers, respectively (Table S2 and S3). All constructs were confirmed by sequencing. Plasmids were expressed in *E. coli* BL21 (DE3) pLysS (Invitrogen). Cells grown in LB broth at 37°C to an OD_600_ of 0.5 were treated with 1mM isopropyl β-D-1-thiogalactopyranoside. After 3 h, the cells were harvested, disrupted by sonication, and centrifuged to obtain cell-free supernatants. 6XHis-tag fusion proteins were purified using Ni-NTA affinity chromatography (Qiagen) as per manufacturer’s instructions. DksA protein was purified as described previously^49^. A GST-DksA fusion protein was purified using Glutathione-Sepharose 4B (bioWORLD, Dublin, OH) according to manufacturer’s protocols. To remove the GST tag, PreScission protease was added to recombinant GST-DksA protein in PBS containing 10 mM DTT. After overnight incubation at 4°C, proteins were eluted and further purified by size-exclusion chromatography on Superdex 75 (GE Healthcare Life Sciences). Purified DksA proteins were aliquoted inside a BACTRON anaerobic chamber (Shel Lab, Cornelius, OR). The purity and mass of the recombinant proteins were assessed by SDS/PAGE.

### Animal studies

Six to eight-week-old immunocompetent C57BL/6, and immunodeficient iNOS^-/-^ or *nox2*^-/-^ mice deficient in the inducible nitric oxide synthetase or the gp91*^phox^* subunit of the NADPH oxidase, respectively, were inoculated intraperitoneally with ∼100 CFU of *Salmonella* grown overnight in LB broth at 37°C in a shaking incubator. Mouse survival was monitored over 14 days. The bacterial burden was quantified in livers and spleens 3 days post infection by plating onto LB agar containing the appropriate antibiotics. Competitive index was calculated as (strain 1/ strain 2)_output_ / (strain 1/ strain 2)_input_. The data are representative of two to three independent experiments. All mice experiments were conducted according to protocols approved by the Institutional Animal Care and Use Committee at the University of Colorado School of Medicine.

### Susceptibility to H_2_O_2_

*Salmonella* strains grown overnight in LB broth at 37°C with shaking were diluted in phosphate buffer saline (PBS) to a final concentration of 2 × 10^5^ CFU/ml. The cells were treated with 200 μM of H_2_O_2_ at 37°C for 2 h. The surviving bacteria were quantified after plating 10-fold serial dilutions onto LB agar. The percent survival was calculated by comparing the surviving bacteria after H_2_O_2_ challenge to the starting number of cells. The effect of H_2_O_2_ on bacterial growth was also examined. Briefly, *Salmonella* strains grown overnight in LB broth at 37°C with shaking were subcultured 1:100 into EG minimal medium. 200 μl were seeded onto honeycomb microplates and treated with or without 400 μM H_2_O_2_. The OD_600_ was recorded every 15 min for up to 40 h in a Bioscreen C plate reader (Growthcurves USA).

### Growth kinetics

Overnight *Salmonella* cultures grown in MOPS-GLC medium with appropriate antibiotics were diluted 1:100 into fresh MOPS-GLC medium. Where indicated, 0.4% D-glucose, 40 μg/ml of each amino acid, 2 mM GSH, 0.4% Casamino acids or 0.15% TCA intermediates was added to MOPS-GLC medium. In addition, MOPS minimal medium was supplemented with 0.4% maltose, fructose, galactose or lactose as needed. *Salmonella* were also grown in E-salts minimal medium supplemented with either 0.4% D-glucose (EG). Bacterial growth was followed by recording OD_600_ values every hour for 7-10 h at 37°C in an aerobic shaking incubator or anaerobic chamber.

### Thin layer chromatography

Nucleotides were examined as originally described with minor modifications^50, 51^. Briefly, *Salmonella* strains grown overnight in MOPS-GLC medium supplemented with 2 mM K_2_HPO_4_ were diluted 1:100 into fresh 0.4 mM K_2_HPO_4_ MOPS-GLC medium. The cultures were grown to early exponential phase till the OD_600_ reached ∼0.2. One-milliliter culture aliquots were labeled with 10 μCi of inorganic ^32^P. After 1h, the cells were treated with 0.4 ml of ice-cold 50% formic acid and incubated on ice for at least 20 min. The extracts were centrifuged at 16,000 × g for 5 min. A 2.5- or 5-μl volume of ice-cold extracts were spotted along the bottom of polyethyleneimine-cellulose TLC plates (Millipore). The spots were air dried, and the TLC plates were placed into a chamber containing 0.9 M K_2_HPO_4_, pH 3.4. ^32^P-labeled nucleotides in the TLC plates were visualized with phosphor screens on a phosphorimager (Bio-Rad), and relative nucleotide levels were quantified with the ImageJ software (NIH).

### Polarographic O_2_ and H_2_O_2_ measurements

Consumption of O_2_ was measured using an ISO-OXY-2 O_2_ sensor attached to an APOLLO 4000 free radical analyzer (World Precision Instruments, Inc., Sarasota, FL) as described^52^. Briefly, 3 ml of *Salmonella* grown aerobically to OD_600_ O.25 in MOPS-GLC medium were rapidly withdrawn, vortexed for one minute and immediately recorded for O_2_ consumption. A two-point calibration for 0 and 21% O_2_ was done as per manufacturer’s instructions. H_2_O_2_ was measured in an APOLLO 4000 free radical analyzer using a H_2_O_2_ specific probe.

### Intrabacterial redox potential

Intrabacterial redox potential was determined by fluorescent measurement for roGFP2 in *Salmonella* as described^53^. The plasmid pFPV25, that encodes and constitutively expresses roGFP2, was electroporated into wild-type *Salmonella* strain 14028s and Δ*greAB* mutant *Salmonella*. Overnight bacterial cultures were subcultured 1:100 in MOPS-GLC medium (without antibiotics) and grown at 37°C with shaking to an OD_600_ of 0.25. Culture aliquots (3 ml) were left untreated or treated with 400 μM H_2_O_2_ for 1 min at 37°C before fluorescence measurement at excitation wavelengths of 405 and 480 nm (roGFP2_ox_ and roGFP2_red_, respectively). Emission was read at 510 nm in both instances. All values were normalized to the ratios obtained for fully oxidized and fully reduced cultures of the respective strains 1 min after treatment with 50 mM H_2_O_2_ or 10 mM DTT, respectively.

### NAD(P)H and NAD(P)^+^ measurements

Intracellular NAD(P)H/NAD(P)^+^ measurements were carried out according as described^52^ with slight modifications. Briefly, *Salmonella* grown in MOPS-GLC medium at 37°C to an OD_600_ of 0.25 were treated for 30 min with or without 400 μM H_2_O_2_. NAD(P)H and NAD(P)^+^ were extracted from pellets in 0.2 M NaOH or 0.2 M HCl. Ten microliter of extracts were added to 90 μl of reaction buffer containing 200 mM bicine, pH 8.0, 8 mM EDTA, 3.2 mM phenazine methosulfate and 0.84 mM 3-(4,5-dimethylthiazol-2-yl)−2,5-diphenyltetrazolium bromide. NAD^+^/NADH and NADP^+^/NADPH concentrations were estimated in reactions containing 20% ethanol and 0.4 μg alcohol dehydrogenase or 2.54 mM glucose-6-phosphate and 0.4 μg glucose-6-phosphate dehydrogenase, respectively. NADH(P) and NAD(P)^+^ measured at 570 nm for the thiazolyl tetrazolium blue cycling assay and calculated by regression analysis of known standards, and specimens were standardized according to OD_600_.

### LC-MS amino acid analysis

To measure amino acids by LC-MS, approximately 5×10^10^ *Salmonella* were collected from cultures grown in MOPS-GLC medium at 37°C to an OD_600_ of 0.25. Amino acids were extracted on ice-cold lysis buffer [5:3:2 ratio of methanol-acetonitrile-water (Fisher Scientific, Pittsburgh, PA)] containing 3 µM of amino acid standards [Cambridge Isotope Laboratories, Inc., Tewksbury, MA]). Samples were vortexed for 30 min at 4°C in the presence of 1-mm glass beads. Insoluble proteins and lipids were pelleted by centrifugation at 12,000 x g for 10 min at 4°C. The supernatants were collected and dried with a SpeedVac concentrator. The pellets resuspended in 0.1% formic acid were analyzed in a Thermo Vanquish ultrahigh-performance liquid chromatography (UHPLC) device coupled online to a Thermo Q Exactive mass spectrometer. The UHPLC-MS methods and data analysis approaches used were described previously^54^.

### Glucose utilization

Glucose in the culture medium was measured by the Glucose Assay Kit (Abcam) as per manufacturer’s instructions. Briefly, *Salmonella* grown in MOPS-GLC medium at 37°C to OD_600_ of 0.25 were harvested and resuspended in fresh MOPS-GLC media. Selected samples were treated with 400 μM H_2_O_2_ at 37°C. Culture supernatants were collected at 2 h after treatment and stored at −20°C until further use. Five microliter of culture supernatants mixed with 400 μl o-toluidine reagent were incubated at 100°C for 8 min. Reaction mixtures were cooled down in ice for 5 min and the OD was recorded at 630 nm. Glucose concentration was calculated by regression analysis of known glucose standard.

### 2-phosphoglycerate and pyruvate estimation

2-phosphoglycerate (2-PG) and pyruvate were measured in *Salmonella* grown in MOPS-GLC medium at 37°C to OD_600_ of 0.25. Where indicated, cells were challenged for 30 min with 400 μM H_2_O_2_. Bacterial cells were harvested and sonicated in 200 μl of ice-cold lysis buffer (25mM Tris-HCl, pH 8.0, 100 mM NaCl). Soluble cytoplasmic extracts obtained after clarification at 13,000 g for 10 min at 4°C were processed with the 2-phosphoglycerate Assay Kit (Abcam) as per manufacturer’s instructions. 2-PG and pyruvate concentrations in the lysates were calculated by linear regression using known 2-PG and pyruvate standards.

### Aldehyde and D-lactate measurement

Aldehyde and D-lactate production in *Salmonella* strains were measured by the Aldehyde Assay Kit and D-Lactate Assay Kit (Sigma-Aldrich), respectively, as per manufacturer’s instructions. Briefly, *Salmonella* strains were grown in MOPS-GLC medium at 37°C to an OD_600_ of 0.25. Because of the growth defect, Δ*greAB* mutant *Salmonella* were grown for extra 2.5 h until they reach an OD_600_ of ∼ 0.25. The harvested cells were sonicated in 200 μl of ice-cold lysis buffer (25mM Tris-HCl; pH 8.0 and 100 mM NaCl). Insoluble material was removed by centrifugation at 13,000 g for 10 min at 4°C. Soluble cytoplasmic extracts were estimated in samples standardized to equal amount of total protein.

### GAPDH enzymatic activity

GAPDH activity in *Salmonella* was measured by the GAPDH activity assay kit (Abcam) as per manufacturer’s instructions. Briefly, *Salmonella* were grown in MOPS-GLC medium at 37°C to an OD_600_ of 0.25. Cells were sonicated in 200 μl of ice-cold lysis buffer (25mM Tris-HCl, pH 8.0,100 mM NaCl). Insoluble material was removed by centrifugation at 13,000 g for 10 min at 4°C. GAPDH enzymatic activity in soluble cytoplasmic extracts was estimated by measuring the accumulation of NADH at 450 nm formed in conversion of glyceraldehyde-3-phosphate into 1, 3-bisphosphate glycerate. GAPDH activity was standardized to equal amounts of protein. GAPDH activity was calculated by linear regression using known NADH standard.

### ATP measurements

ATP concentrations were quantified with the luciferase-based ATP determination kit (Molecular Probes). Briefly, *Salmonella* grown in MOPS-GLC medium at 37°C to an OD_600_ of 0.25 were challenged for 30 min with and without 400 μM H_2_O_2_. Pellets from 2 ml cultures were thoroughly mixed with 0.5 ml of ice-cold, 380 mM formic acid containing 17 mM EDTA. After centrifugation for 1 min at 16,000×g, supernatants were diluted 25-fold into 100 mM N-tris(hydroxymethyl)methyl-2-aminoethanesulfonic acid (TES) buffer, pH 7.4. Ten-microliter of samples or ATP standards were mixed with 90 μl of reaction master mix (8.9 ml of water, 500 μl of 20× buffer, 500 μl of 10 mM D-luciferin, 100 μl of 100 mM dithiothreitol [DTT], 2.5 μl of 5-mg/ml firefly luciferase). Luminescence was recorded in an Infinite 200 PRO (Tecan Life Sciences). ATP concentrations were calculated by linear regression using ATP standards, and the intracellular concentration of ATP was standardized to CFU/ml.

### RNA isolation, library preparation and RNA seq

*Salmonella* grown in MOPS-GLC medium at 37°C to an OD_600_ of 0.25 were treated with 1 ml of TRIzol reagent (Life Technologies). Following chloroform extraction, RNA was precipitated from the aqueous phase by the addition of 3 M sodium acetate (1/10, vol/vol), 50 mg/ml glycogen (1/50, vol/vol), and an equal volume of 100% isopropyl alcohol. Precipitated RNA was washed twice with 70% (vol/vol) ethanol, suspended in RNase free dH_2_O, and treated with RNase free DNase I, according to the supplier’s specifications (Promega). Reactions were terminated by the addition of an equal volume of phenol/chloroform/ isoamyl alcohol solution (25:24:1) (PCI). The aqueous phase was treated with an equal volume of chloroform. RNA in the resulting aqueous phase was precipitated by the addition 3 M sodium acetate (1/10 vol/vol), 50 mg/mL glycogen (1/50 vol/vol and 3 volumes of 100% ethanol. The quality of the isolated RNA was assessed on an Agilent Bioanalyzer. Ribosomal RNA was removed from the total RNA preparation using the MICROBExpress kit (Life Technologies). Starting with 1 μg purified mRNA, samples were fragmented with the NEB Magnesium Fragmentation module at 94°C for 5 min. RNA was purified by PCI extraction and ethanol precipitation and sodium acetate, and libraries were prepared for Illumina sequencing by following the protocol accompanying the NEBNext Ultra RNA Library Prep Kit through completion of the second strand synthesis step. Libraries were made by NEBNext Ultra RNA Library Prep Kit protocol for a target insert size of 300 bp. Samples were barcoded using NEBNext Multiplex Oligos (Universal primer, Index Primers Set 1 and Index Primers Set 2), and the resulting indexed libraries were sequenced on an Illumina MiSeq using 300-nt reads. The i7 Illumina adapters were trimmed from raw paired reads by utilizing Cutadapt version 2.10 in the Linux terminal with the sequences AGATCGGAAGAGCACACGTCTGAACTCCAGTCAC and AGATCGGAAGAGCGTCGTGTAGGGAAAGAGTGTAGATCTCGGTGGTCGCCGTATCATT for the forward and reverse reads, respectively. Reads were then mapped with Bowtie2 ^55, 56^ version 2.3.2 using CP001363.1 and CP001362.1^57^ as the reference genome for *S. Typhimurium 14028s*. Picard version 2.18.27 was then used to remove duplicates and sort the reads. HTseq^58^ version 0.13.5 was then leveraged to generate count files by locus for each sample. Counts for each sample were then statistically analyzed utilizing DEseq2 1.30.1^59^ and edgeR 3.32.1^60, 61^ in R Studio running R version 4.0.4 by using Fisher’s exact test on the tagwise dispersion of counts for loci that had at least 80 reads total across all samples be analyzed. Genes categorized following KEGG annotations were imported with heatmap 1.0.12 in R for graphical representation along the following color breaks for fold-change values of: 0.1386, 0.6060, 1.0000, 2.1304, 2.8125, and 7.5938. Volcano plots were generated with EnhancedVolcano in R. PCA analysis was performed after a log transformation and Pareto scale of the raw counts data. Final heatmaps, PCA and loadings graphs were manipulated in Inkscape version 0.92.1 to add labels and overlay findings.

### RNA isolation and quantitative RT-PCR

*Salmonella* strains grown in MOPS-GLC medium in a shaking incubator at 37°C to an OD_600_ of 0.25 were centrifuged at 16,000 × g for 10 min at 4°C. The bacterial pellets were saved at −80°C until further processing. DNA-free RNA was purified using a High Pure RNA isolation kit (Roche) according to the manufacturer’s instructions. First-strand cDNA generation from total RNA was generated using Moloney murine leukemia virus (M-MLV) reverse transcriptase (Promega). Relative mRNA quantitation was done using the SYBR green quantitative real-time PCR (qRT-PCR) master mix (Roche) using the primers described in Table S3. Data evaluation of 3 biological replicates done in triplicate was performed using the threshold cycle (2^-ΔΔCT^) method. Gene expression was normalized to internal levels of the housekeeping gene *rpoD*. Transcripts that exhibited 2-fold up- or downregulation were considered to exhibit a significant change.

### *In vitro* transcription

Products synthesized in *in vitro* transcription reactions were quantified by qRT-PCR as described previously^27, 49^. Transcription reactions were performed in 40 mM HEPES, pH 7.4, 2 mM MgCl_2_, 60 mM potassium glutamate, 0.1% Nonidet P-40, 200 μM of each ATP, GTP, CTP and UTP (Thermo Fisher Scientific, Grand Island, NY), 8 U RiboLock RNase inhibitor (Thermo Fisher Scientific, Grand Island, NY), 1 nM of DNA template, 5 nM *E. coli* holoenzyme RNA polymerase (New England Biolabs, Ipswich, MA). Where indicated, 0-200 nM of GreA or 0-100 nM of GreB proteins were added to the *in vitro* transcription reactions. Reactions were incubated at 37°C for 10 min, and then heat-inactivated at 70°C for 10 min. After DNase I treatment, template DNA was removed by DNA-free DNA Removal kit (Thermo Fisher Scientific, Grand Island, NY). The resulting materials were used as template to generate cDNA using 100 U Moloney murine leukemia virus (MMLV) reverse transcriptase (Promega, Madison, WI), 0.45 μM N6 random hexamer primers (Thermo Fisher Scientific, Grand Island, NY) and 20 U RNase inhibitor (Promega, Madison, WI). The cDNA was synthesized by incubating reaction at 42°C for 30 mins. Relative mRNA quantitation was done using the SYBR green quantitative real-time PCR (qRT-PCR) master mix (Roche, Basel, Switzerland) using specific primers (Table S3). Data evaluation of 3 biological replicates done in duplicates or triplicate was performed using the threshold cycle (2^-ΔΔCT^) method.

### Transcriptional pausing

Transcriptional pause assays were performed using PCR-amplified 300 bp promoter with 50 bp coding sequence of either *gapA*, *eno* or *cydA* gene containing *rrnB* and *rpoC* terminator at the 3’-end. Transcriptional forks were initiated in standard transcription buffer (40 mM HEPES, pH 7.4, 2 mM MgCl_2_, 60 mM potassium glutamate, 0.1% Nonidet P-40) containing 8 U RiboLock RNase inhibitor, 10 nM RNAP holoenzyme, 2 nM template DNA with or without 200 nM of GreA, 100 nM GreB, or 1 μM DksA proteins. The reactions were carried out for 9 min at 37°C. Multiround runoff transcription assays were started upon the addition of 200 μM NTPs containing 0.2 μCi [^32^P]-α-UTP (3,000 Ci/mmol). Aliquots removed at between 1 and 9 min were treated with 125 μl of transcription stop buffer (0.6 M Tris-HCl, pH - 8.0 and 20 mM EDTA, pH - 8.0) containing 5 μg tRNA. Samples were precipitated with 3 volumes of 100% ethanol, followed by centrifugation at 12000 rpm for 20 min. The RNA products dissolved in 2X formamide RNA sample buffer were separated in 7 M urea-16% PAGE gels. Transcriptional pause products were identified by using ^32^P-labeled Decade™ Markers System (Ambion) and visualized by the Typhoon PhosphorImager (GE Healthcare).

### Statistical analysis

Statistical analyses were performed using GraphPad Prism 5.0 software. One-way and two-way ANOVA, *t*-tests and logrank tests were used. Data were considered statistically different when *p* < 0.05.

## ACKNOWLEDGMENTS

We thank Dr. Jessica Jones-Carson for kindly providing the mice, and members of the Vazquez-Torres lab for critically reading the manuscript. We also thank Ted R. Shade from the Genomics Shared Resource Facility, University of Colorado Anschutz Medical Campus, for sequencing of the RNA Seq library, Michael Armstrong Mass Spectrometry Facility, Skaggs School of Pharmacy and Pharmaceutical Sciences, University of Colorado Denver – Anschutz Medical Campus, for analysis of amino acids, and Dr. Tonya Brunetti at the Department of Immunology and Microbiology for her guidance representing and preparing sequencing data sets for publication.

## AUTHOR CONTRIBUTIONS

SK, AVT designed and wrote the study; SK, LL performed investigations; JT analyzed data; JSK, provided materials and methodologies; AVT provided funding.

## Supplementary Materials

**Table S1.**
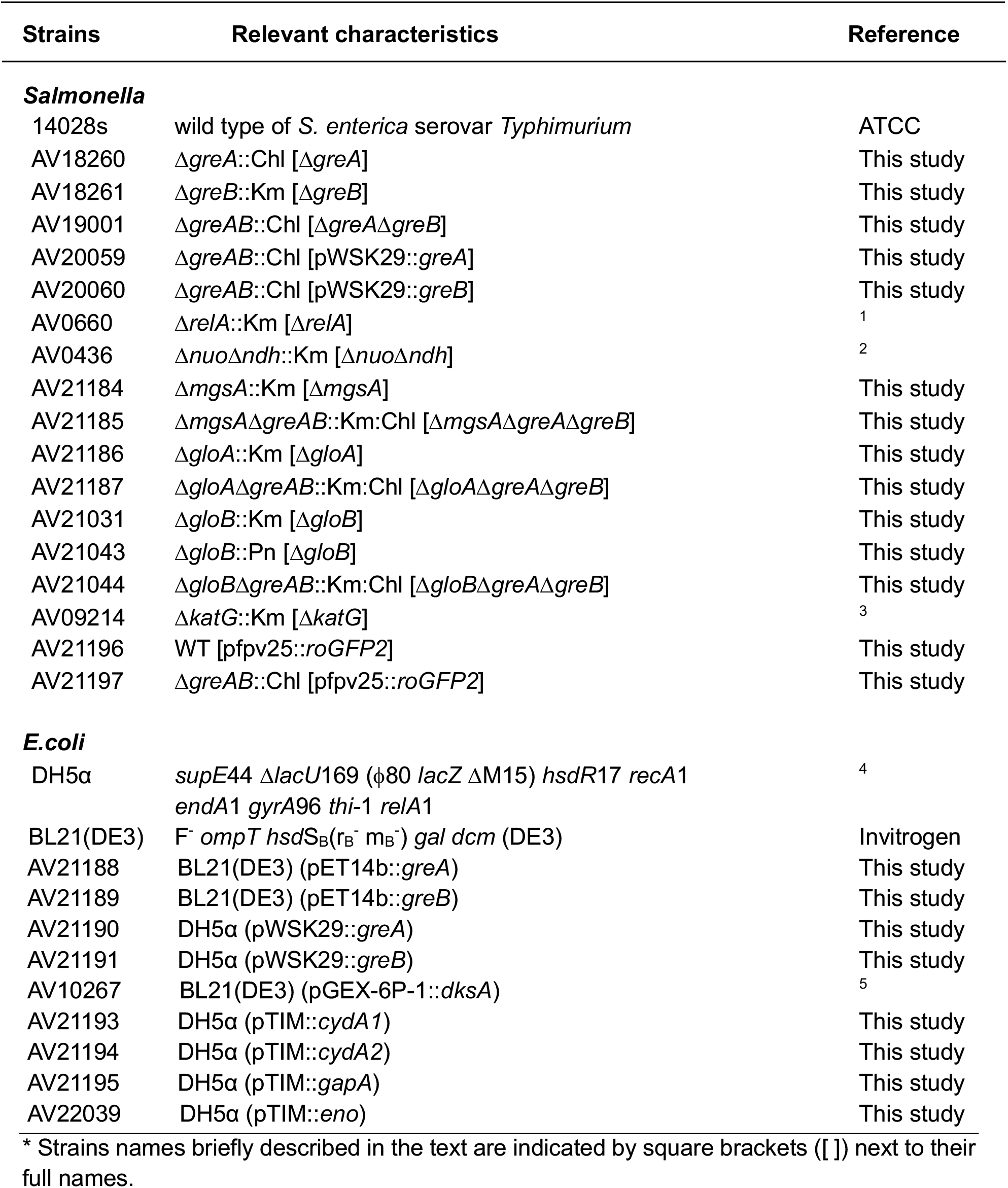
Bacteria used in this study.

**Table S2.** Plasmids used in this study.

| **Plasmid** | **Relevant characteristics** | **Source** |
| --- | --- | --- |
| pET14b | *ori* pBR322, N-terminal 6His-Tag expression vector, Pn<sup>r</sup> | Novagen |
| pTIM | *in vitro* transcription backbone plasmid, bla *rrnB* & *rpoC* term pBluescript, Pn<sup>r</sup> | 6 |
| proGFP2 | pfpv25-roGFP2 | 7 |
| pWSK29 | low copy plasmid, *lacZ*α, Pn<sup>r</sup> | 8 |
| pET14b::*greA* | pET14b + 477 bp *greA* CDS, Pn<sup>r</sup> | This study |
| pET14b::*greB* | pET14b + 474 bp *greB* CDS, Pn<sup>r</sup> | This study |
| pTIM::*cydA* 1 | pTIM + 300 bp *pcydA* and 50 bp *cydA* CDS, Pn<sup>r</sup> | This study |
| pTIM::*cydA* 2 | pTIM + 300 bp *pcydA* and 400 bp *cydA* CDS, Pn<sup>r</sup> | This study |
| pTIM::*gapA* | pTIM + 300 bp *pgapA* and 50 bp *gapA* CDS, Pn<sup>r</sup> | This study |
| pTIM::*eno* | pTIM + 300 bp *peno* and 50 bp *eno* CDS, Pn<sup>r</sup> | This study |
| pWSK29::*greA* | pWSK29 + 877 bp *greA* CDS with promoter, Pn<sup>r</sup> | This study |
| pWSK29::*greB* | pWSK29 + 874 bp *greB* CDS with promoter, Pn<sup>r</sup> | This study |

**Table S3.**
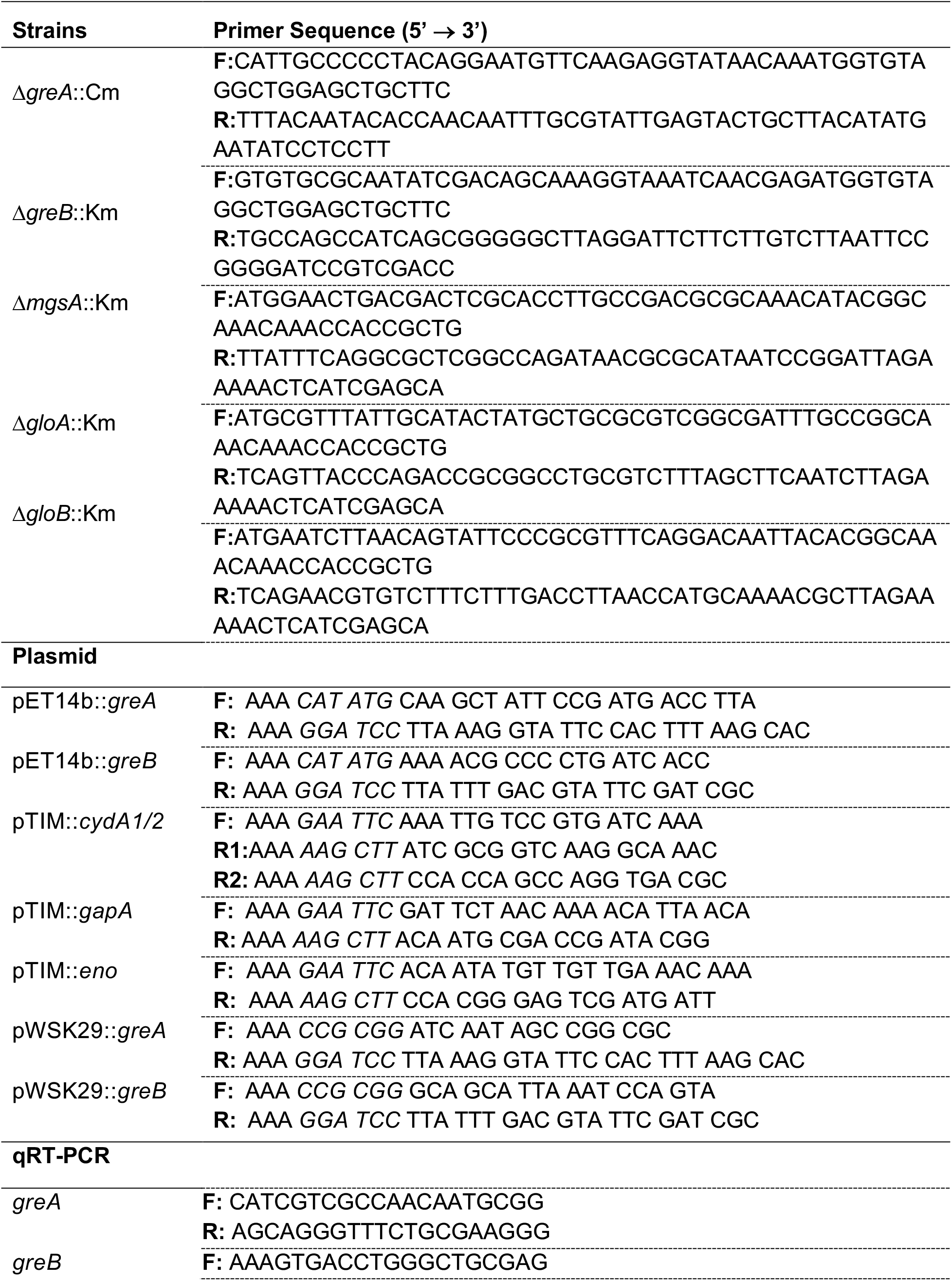
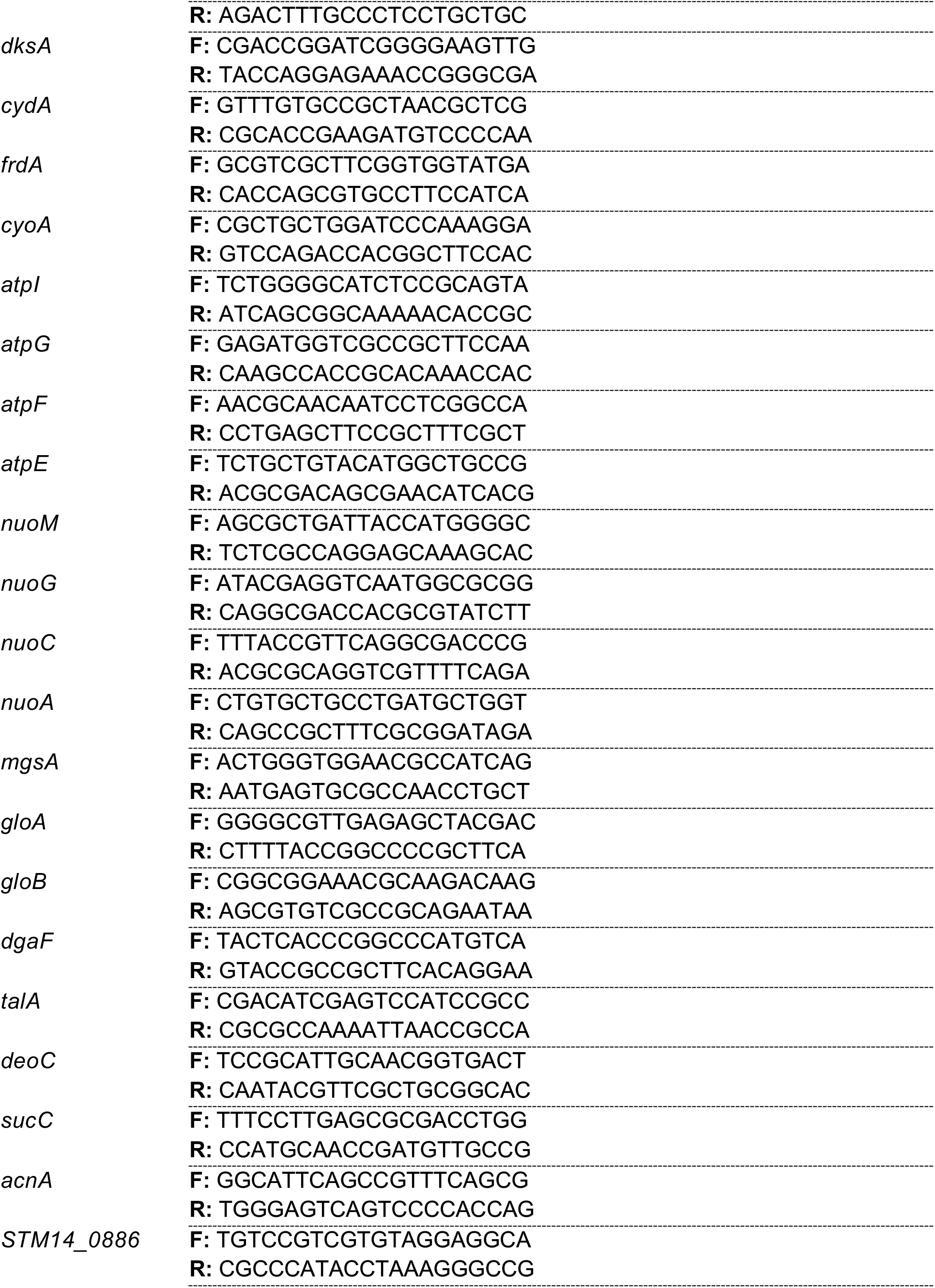
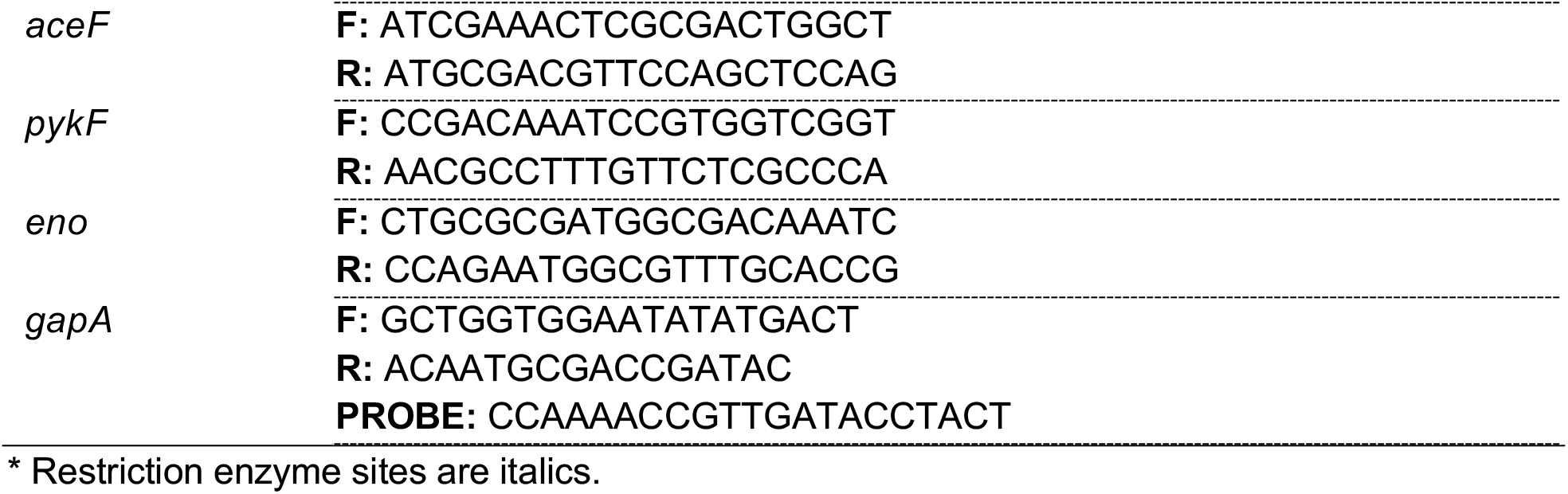
Oligonucleotides used in this study.

### Supplementary figure

**Fig. S1.**
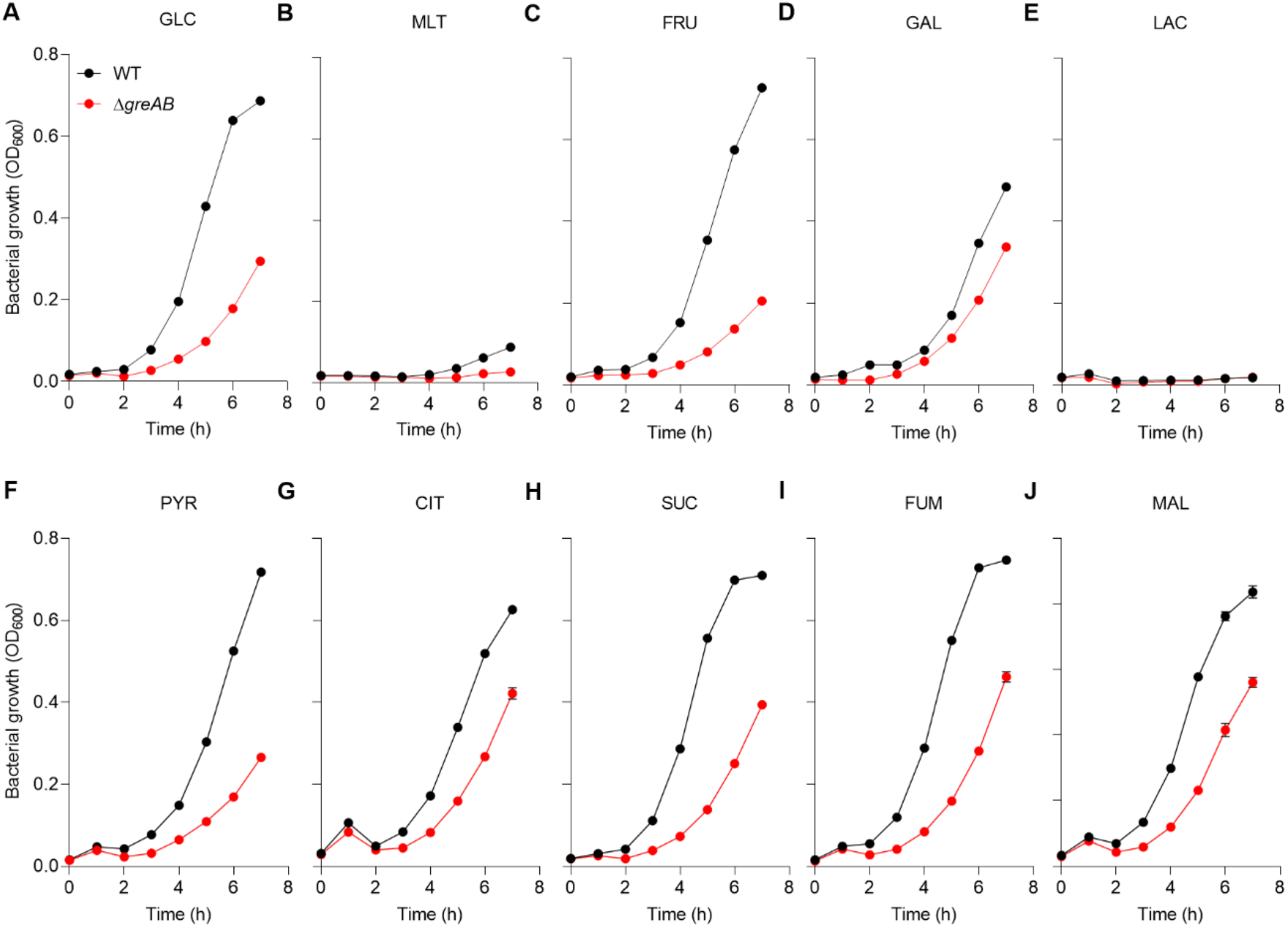
Effect of carbon source on *Salmonella* growth. Growth of wild-type (WT) and !ι*greAB Salmonella* in MOPS minimal media supplemented with 0.4% glucose (A), maltose (B), fructose (C), galactose (D), and lactose (E), or 0.15% pyruvate (F), citrate (G), succinate (H), fumarate (I), or malate (J). Data are shown as mean ± S.D. (N=3). GLC, Glucose; MLT, Maltose; FRU, Fructose; GAL, Galactose; LAC, Lactose; PYR, Pyruvate; CIT, Citrate; SUC, Succinate; FUM, Fumarate; MAL, Malate.

**Fig. S2.**
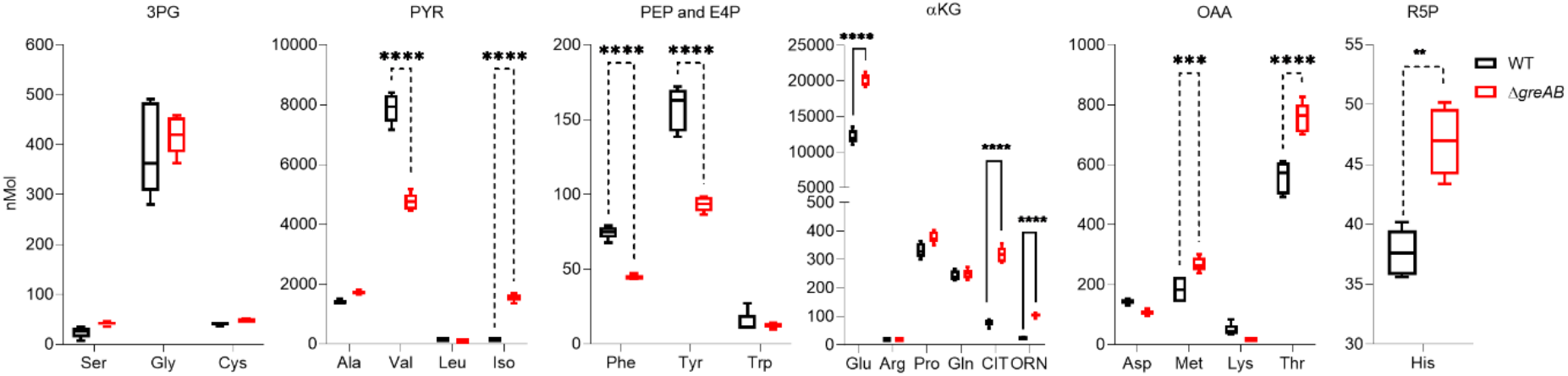
Amino acid pools in *Salmonella* grown on glucose. Amino acids were quantified in *Salmonella* grown in MOPS-GLC minimal medium to an OD_600_ of 0.25 by liquid chromatography mass spectrometry (LC-MS). Data are the mean ± S.D (N=5). ***,****; *p*< 0.001 and *p*< 0.0001, respectively as determined by two-way ANOVA. Three letter amino acid code is used. 3PG, 3 phosphoglycerate; PYR, pyruvate; PEP, phosphoenolpyruvate; E4P, erythrose 4-phosphate; αKG, alpha-ketoglutarate; OAA, oxaloacetic Acid; R5P, ribose 5-phosphate; CIT, citrulline; ORN, ornithine.

**Fig. S3.**
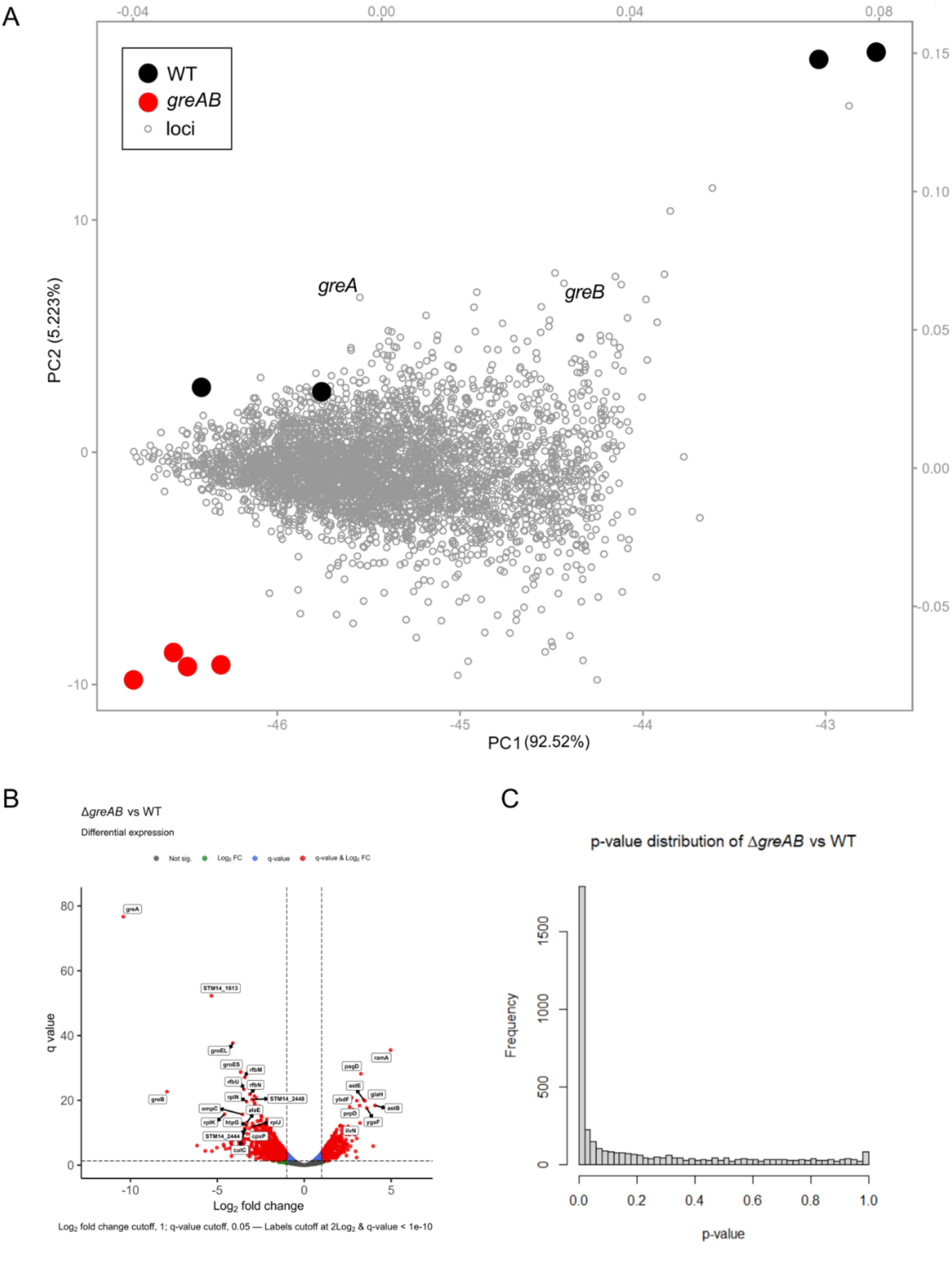
RNA seq analysis of *Salmonella* grown in glucose. (A) A principle-component analysis of RNAseq data obtained from WT and Δ*greAB Salmonella*. The PCA was performed in R on log-transformed and Pareto-scaled counts from each loci. Loadings are shown by the small gray points and with the gray axes. (B) Volcano plot and q-value distributions for comparisons of RNA-seq count data after analysis with DESeq2 and edgeR with tagwise dispersion and fdr-corrected p-values. Data points are labeled where the meet the fold-change and q-value cutoff criteria stated at the bottom of each figure.

**Fig. S4.**
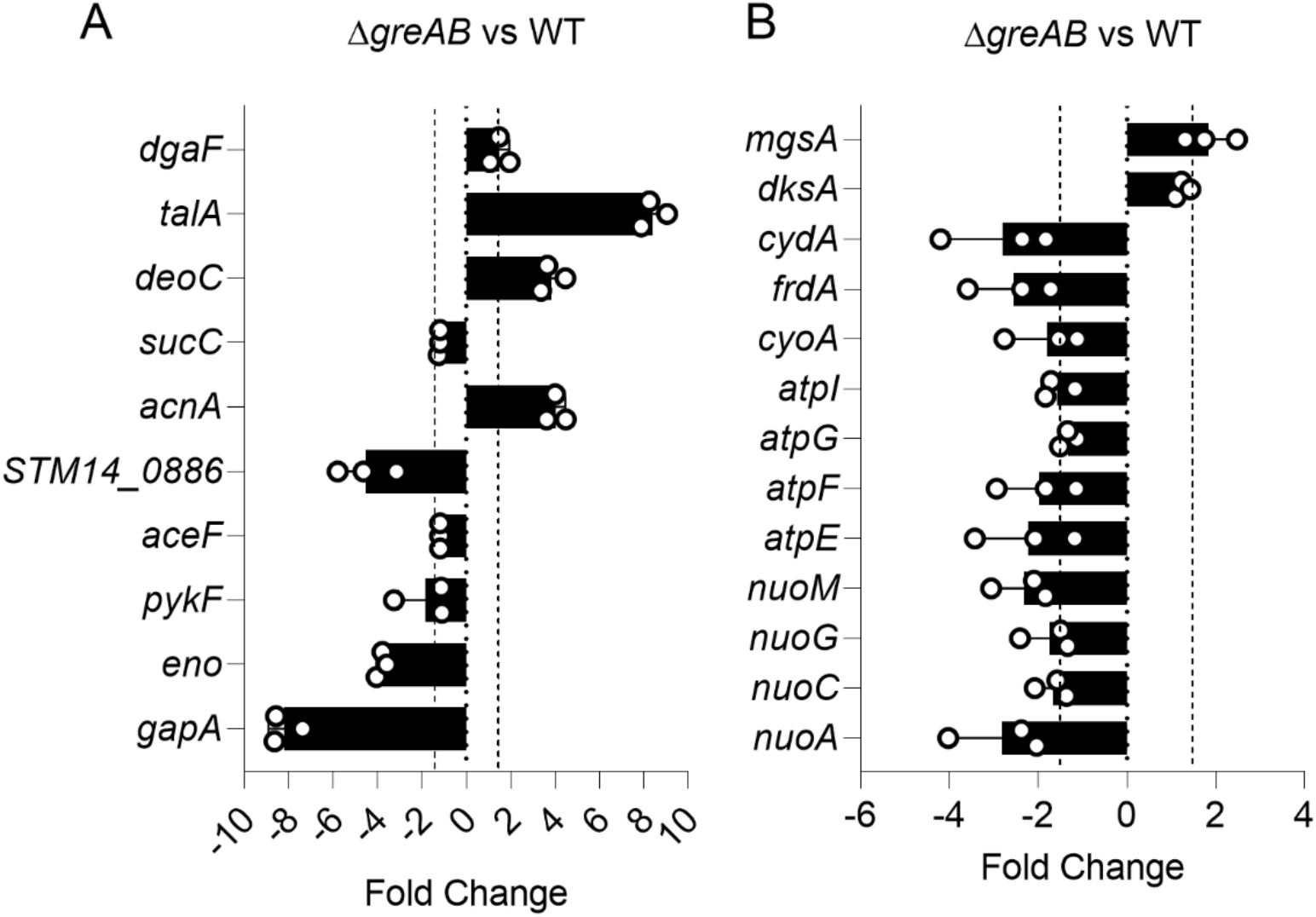
Gene expression in *Salmonella* grown on glucose. qRT-PCR analysis of specimens isolated from WT or !1*greAB Salmonella* grown to an OD_600_ of 0.25 in MOPS-GLC. The data, which are shown in the form of down-regulated and up-regulated black solid bars, represent the average fold-change ± S.D. (N=3). The data were normalized to internal *rpoD* levels. The 1.5-fold up- or downregulation considered to exhibit a significant change.

**Fig. S5.**
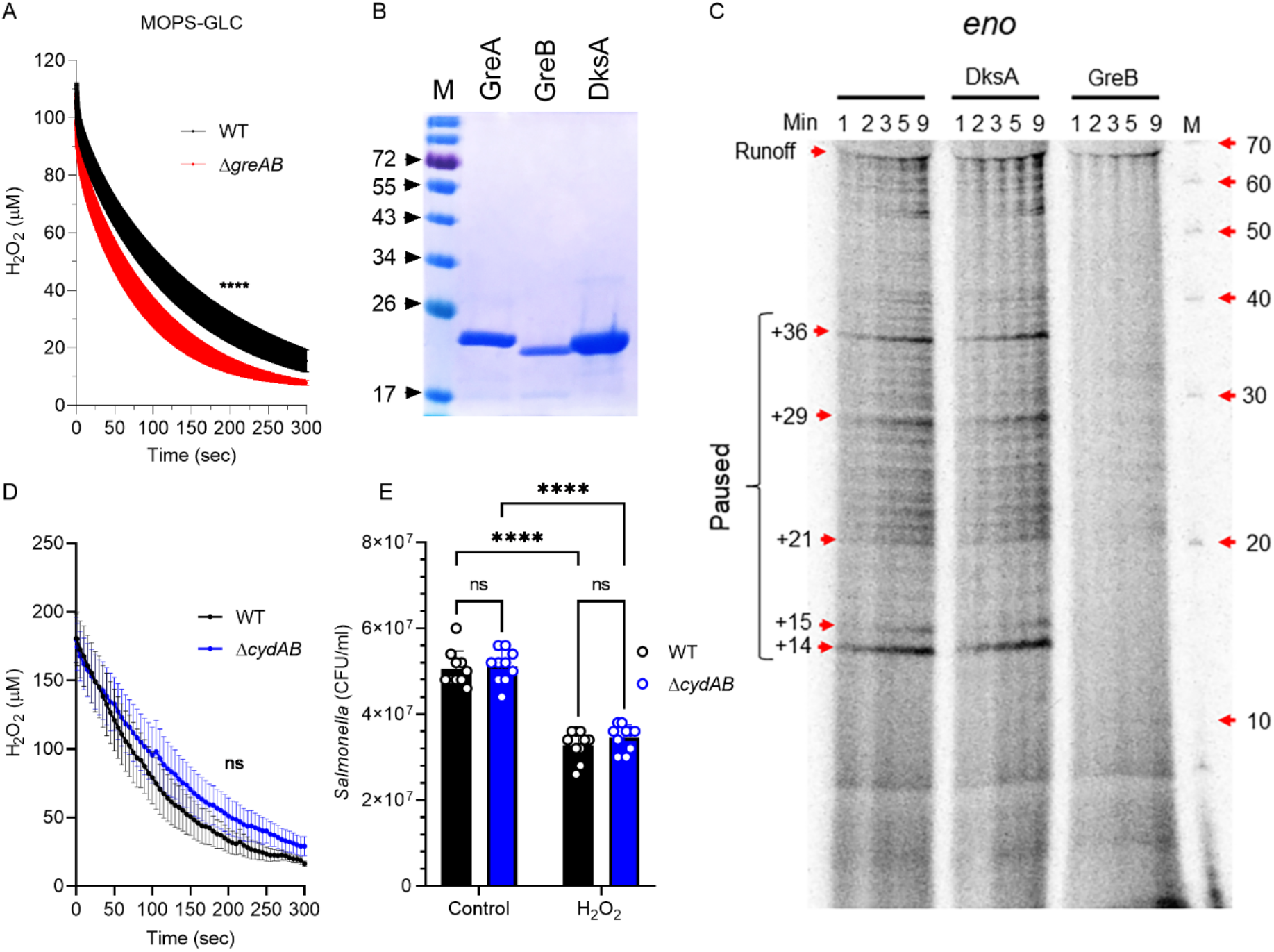
H_2_O_2_ consumption by *Salmonella* and transcriptional pausing. (A) Degradation of H_2_O_2_ by wildtype and !1*greAB Salmonella* grown in MOPS-GLC to an OD_600_ of 0.25. The degradation of H_2_O_2_ was followed polarographycally in an ISO-OXY/HPO analyzer equipped with a H_2_O_2_ probe. At the time of measurement, 100 μM H_2_O_2_ was added to the cultures. Data are shown as mean ± S.D. (N=6). **** *p*< 0.0001 as determined by *t*-test. (B) Purified proteins GreA, GreB and DksA were evaluated by SDS-PAGE gels and visualized by Coomassie Brilliant Blue staining. (C) Transcriptional pausing of *in vitro* transcription reactions containing *eno* templates. Where indicated, the reactions contained 1 μM DksA or 100 nM GreB recombinant proteins. The α^32^P-UTP-labaled products of the *in vitro* transcription reactions were visualized in urea PAGE gels. (D) Degradation of H_2_O_2_ by Δ*cydAB Salmonella* grown to an OD_600_ of 0.25 as measured polarographycally in an ISO-OXY/HPO analyzer equipped with a H_2_O_2_ probe. 200 μM H_2_O_2_ were added to the cultures immediately before measurements were initiated. Data are the mean ± S.D. (N=6). *p*< 0.1116 as determined by *t*-test. (E) CFU of wild-type (WT) and !1*cydAB* mutant *Salmonella* grown for 30 min in MOPS-GLC in the presence or absence of 400 μM H_2_O_2_. Data are shown as mean ± S.D. (N=10). ****, *p*<0.0001 as determined by two-way ANOVA.

